# The SUMO ligase Su(var)2-10 links piRNA-guided target recognition to chromatin silencing

**DOI:** 10.1101/533091

**Authors:** Maria Ninova, Yung-Chia Ariel Chen, Baira Godneeva, Alicia K. Rogers, Yicheng Luo, Alexei A. Aravin, Katalin Fejes Tóth

## Abstract

Regulation of transcription is the main mechanism responsible for precise control of gene expression. While the majority of transcriptional regulation is mediated by a multitude of DNA-binding transcription factors that bind to regulatory gene regions, an elegant alternative strategy employs small RNA guides, piwi-interacting RNAs (piRNAs) to identify targets of transcriptional repression. Here we show that in *Drosophila* the small ubiquitin-like protein SUMO and the SUMO E3 ligase *Su(var)2-10* are required for piRNA-guided deposition of repressive chromatin marks and transcriptional silencing of piRNA targets. Su(var)2-10 links the piRNA-guided target recognition complex to the silencing effector by binding the piRNA/Piwi complex and inducing SUMO-dependent recruitment of the SetDB1/Wde histone methyltransferase effector. We propose that in *Drosophila*, the nuclear piRNA pathway has co-opted a conserved mechanism of SUMO-dependent recruitment of the SetDB1/Wde chromatin modifier to confer repression of genomic parasites.

**Highlights:**

- piRNA-induced transposon silencing requires SUMO and the SUMO E3 ligase Su(var)2-10
- Su(var)2-10 links the target recognition complex to the silencing effector
- Su(var)2-10 binds the piRNA-guided target recognition complex and deposits SUMO on target chromatin
- Su(var)2-10 induces SUMO-dependent recruitment of the SetDB1/Wde histone methyltransferase to target loci

## Introduction

Transcriptional control is the first and most important step in regulating gene expression. The majority of transcriptional control is achieved by transcription factors that bind short sequence motifs on DNA. In many eukaryotic organisms, transcriptional repression can also be guided by small RNAs, which - in complex with Argonaute family proteins - recognize their genomic targets using complementary interactions with nascent RNA transcripts(Holoch and Moazed, 2015). This mode of regulation provides flexibility in target selection without the need for novel transcription factors. Accordingly, this mechanism is well suited for genome surveillance systems to identify and repress the activity of harmful genetic elements such as transposons.

Transcriptional repression guided by small RNAs and Argonautes correlates with the deposition of repressive chromatin marks, particularly histone 3 lysine 9 methylation (H3K9me) in S. *pombe*, plants and animals(Bernatavichute et al., 2008; Enke et al., 2011; Gu et al., 2012; LeThomas et al., 2013; Pezic et al., 2014; Volpe et al., 2002). In addition to H3K9me, plants and mammals also employ CpG DNA methylation for target silencing(Aravin et al., 2008; Mette et al., 2000). The molecular mechanism of small RNA/Ago induced transcriptional gene silencing is best understood in S. *pombe*, where the RNA-induced transcriptional silencing complex (RITS) was studied biochemically and genetically(Holoch and Moazed, 2015; Verdel et al., 2004). In contrast to yeast, the molecular mechanism of RNA-induced transcriptional silencing in higher eukaryotes remains poorly understood. Importantly, mechanisms of small RNA-induced transcriptional repression might have independently evolved several times during evolution. Thus, RITS-mediated silencing in Metazoa might mechanistically differ from that of S. *pombe.*

In Metazoa, small RNA guided transcriptional repression is mediated by Piwi proteins, a distinct clade of the Argonaute family, and their associated Piwi-interacting RNAs (piRNAs). Both in *Drosophila* and in mouse, the two best-studied metazoan systems, nuclear Piwis are responsible for transcriptional silencing of transposons, which ensures the first line of defence against their expression(Aravin et al., 2008; Carmell et al., 2007; Kuramochi-Miyagawa et al., 2008; LeThomas et al., 2013; Manakov et al., 2015; Pezic et al., 2014; Rozhkov et al., 2013; Sienski et al., 2012). The current model proposes that target recognition is achieved through binding of the Piwi/piRNA complex to nascent transcripts of target genes. In both *Drosophila* and mouse, piRNA-dependent silencing of transposons correlates with accumulation of repressive chromatin marks (H3K9me3 and, in mouse, CpG methylation of DNA) on target sequences(Carmell et al., 2007; Kuramochi-Miyagawa et al., 2008; LeThomas et al., 2013; Pezic et al., 2014; Rozhkov et al., 2013; Sienski et al., 2012). Both H3K9me3 and CpG methylation are well-known repressive marks that can recruit repressor proteins, such as HP1a(Maison and Almouzni, 2004), providing a mechanism for transcriptional silencing. However, how recognition of nascent RNA by the Piwi/piRNA complex leads to deposition of repressive marks at the target locus is not well understood. Piwi proteins lack domains that might induce chromatin modifications and transcriptional silencing. Genetic screens in *Drosophila* identified two proteins, Asterix (Arx)/Gtsf1 and Panoramix (Panx)/Silencio, which are required for transcriptional silencing and which physically associate with Piwi(Donertas et al., 2013; Muerdter et al., 2013; Ohtani et al., 2013; Sienski et al., 2015; Yu et al., 2015). Accumulation of the repressive H3K9me3 mark on targets of Piwi and Panx were shown to require the activity of the histone methyltransferase SetDB1 (also known as Egg)(Rangan et al., 2011; Sienski et al., 2015; Yu et al., 2015), however, a mechanistic link between the Piwi/Arx/Panx complex, which recognizes targets, and the effector chromatin-modifier has not been established.

Here we identified Su(var)2-10 protein as a novel player in piRNA-induced transcriptional silencing that provides the link between the Piwi/piRNA and the SetDB1 complex. In *Drosophila, Su(var)2-10* mutation causes suppression of position effect variegation, a phenotype indicative of its involvement in chromatin repression(Elgin and Reuter, 2013; Reuter and Wolff, 1981). Su(var)2-10 was shown to associate with chromatin and regulate chromosome structure(Hari et al., 2001) and also emerged in several screens as a putative interactor of the central heterochromatin component HP1 and a repressor of enhancer function (Alekseyenko et al., 2014; Stampfel et al., 2015; Stielow et al., 2008a), however its molecular functions in chromatin silencing were not investigated. Su(var)2-10 belongs to conserved PIAS/Siz protein family(Betz et al., 2001; Hari et al., 2001; Mohr and Boswell, 1999), of which the yeast and mammalian homologs act as E3 ligases for SUMOylation of several substrates(Johnson and Gupta, 2001; Kahyo et al., 2001; Kotaja et al., 2002; Sachdev et al., 2001; Schmidt and Muller, 2002; Takahashi et al., 2001). We studied the role of Su(var)2-10 in germ cells of the ovary, where chromatin maintenance and transposon repression is essential to grant genomic stability across generations. We found that Su(var)2-10 depletion from germ cells phenocopies loss of the nuclear Piwi protein: both lead to strong transcriptional activation of transposons, and loss of repressive chromatin marks over transposon sequences. Su(var)2-10 genetically and physically interacts with Piwi and its auxiliary factors, Arx and Panx, and is required for Panx-induced transcriptional silencing. Recruitment of Su(var)2-10 to a target locus induces local transcriptional repression and H3K9me3 deposition via the methyltransferase SetDB1. Finally, we demonstrate that the repressive function of Su(var)2-10 is dependent on its SUMO E3 ligase activity and the SUMO pathway. Tethering of Su(var)2-10 to chromatin induces local accumulation of SUMO, which can directly interact with the conserved SetDB1 co-factor Wde (MCAF1/ATF7IP in mammals). Together, our data points to a model in which Su(var)2-10 acts downstream the piRNA/Piwi complex to deposit local SUMOylation, which in turn leads to the recruitment of the silencing SetDB1/Wde complex. SUMO modification was shown to play a role in the formation of silencing chromatin in various systems from yeast to mammals, including the recruitment of the silencing effector SETDB1 and its co-factor MCAF1 by repressive transcription factors (Ivanov et al., 2007; Maison et al., 2011, 2016; Shin et al., 2005; Stielow et al., 2008b; Thompson et al., 2015; Uchimura et al., 2006). Together, these findings indicate that the piRNA pathway utilizes a conserved mechanism of silencing complex recruitment through SUMOylation to confer transcriptional repression.

## Results

### Germline-specific knockdown of *Su(var)2-10* induces embryonic lethality

The essential *Su(var)2-10* gene encodes a conserved protein that is highly expressed in the *Drosophila* ovary (FlyAtlas(Chintapalli et al., 2007)). To investigate the role of *Su(var)2-10* in the germline, we employed germline-specific knockdown (GLKD) in the *Drosophila* ovary using two different short hairpin RNAs (shRNAs) that target the *Su(var)2-10* mRNA, shSv210-1 and shSv210-2, driven by the maternal tubulin-GAL4. Expression of each shRNA resulted in ∼95% reduction of the Su(var)2-10 mRNA in the ovary (Fig. 1A). The level of ectopically expressed GFP-tagged Su(var)2-10 protein was also reduced by both shRNAs, with shSv210-2 showing stronger depletion than shSv210-1 (Fig. 1B).

**Figure 1.**
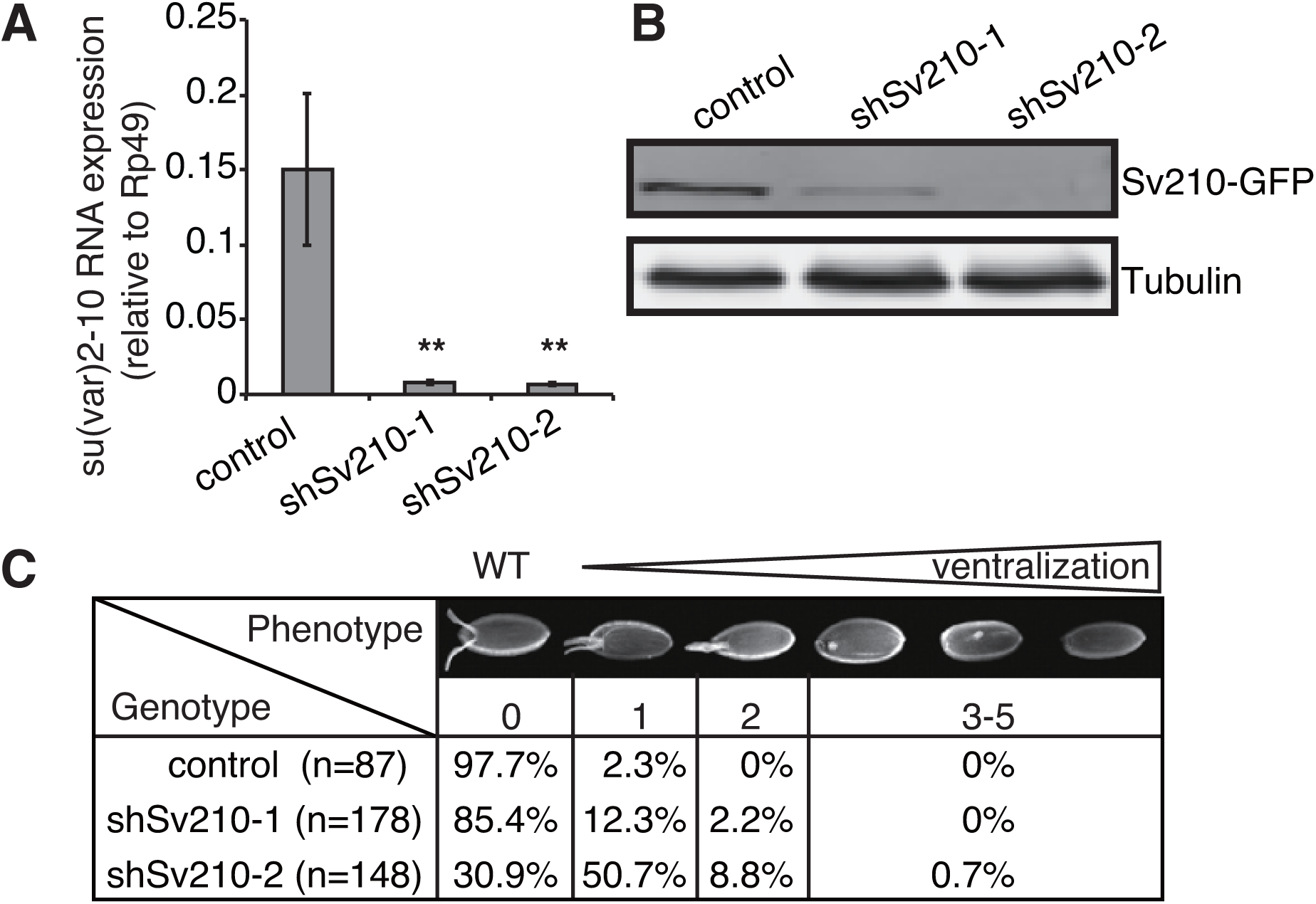
Su(var)2-10 depletion in the *Drosophila* female germline leads to embryo ventralization. (A)Knockdown of *Su(var)2-10* using two different shRNAs (shSv210-1 and shSv210-2) leads to reduced transcript level. Bar plots show the relative levels of Su(var)2-10 mRNA in shW, shSv210-1 and shSv210-2 ovaries as estimated by RT-qPCR. *Rp49* mRNA was used as an endogenous control; error bars show the standard deviation from three biological replicas. Statistical significance was determined by two-tailed Student’s t-test. **p<0.01 (B)Su(var)2-10 protein level is reduced upon knockdown. Western blot shows the levels of GFP-tagged Su(var)2-10 protein expressed in ovaries of control (shW) and Su(var)2-10 knockdown flies (shSv210-1 and shSv210-2), determined by anti-GFP antibody. Tubulin (Tub) is used as loading control. (C)Depletion of Su(var)2-10 causes egg shell ventralization. Top panel illustrates different classes of ventralization phenotypes ordered by severity (Images adopted from Meignin and Davis(Meignin and Davis, 2008)) Table shows the proportion of eggs from control (shW) and Su(var)2-10 knockdown ovaries displaying each phenotype. Both shRNAs targeting Su(var)2-10 led to partial fusion of dorsal appendages.

Next we examined the effect of Su(var)2-10 knockdown on germline development. Ovaries from females with GLKD of Su(var)2-10 did not show gross phenotypic difference in ovarian morphology. Such females produce eggs, but no viable progeny, indicating that Su(var)2-10 plays an important role during gametogenesis. Embryos produced by females with GLKD of Su(var)2-10 showed a varying degree of ventralization phenotype, indicating defects in axis specification (Fig. 1C). Consistent with the stronger protein depletion, shSv210-2 had a stronger penetrance, with 60% of the embryos showing mild to severe ventralization.

We also addressed Su(var)2-10 function in the somatic follicular cells of the ovary, using shRNA under the control of traffic jam (Tj) GAL4. Depletion of Su(var)2-10 in follicular cells caused severe morphological defects and collapse of oogenesis phenocopying effects Piwi depletion in follicular cells. Together, these data indicate that Su(var)2-10 plays an important role in both germ cells and the ovarian somatic cells that supports germline development.

### Depletion of Su(var)2-10 induces transposon derepression

Embryonic ventralization is a known consequence of DNA damage in the ovary(Abdu et al., 2002; Ghabrial et al., 1998) that can be induced by several mechanisms including activation of endogenous transposable elements(Klattenhoff et al., 2007). Measuring expression of several transposon families by RT-qPCR showed that the germline-specific transposon HetA is strongly upregulated in ovaries of both Su(var)2-10 GLKD lines (∼15 and ∼150 fold) (Fig. 2A). In contrast, Blood, a transposon expressed in both ovarian germline and soma, and ZAM, a transposon restricted to the somatic cells in the ovary, were not significantly affected. We analysed global changes in transposon and gene expression upon GLKD of Su(var)2-10 by RNA sequencing (RNA-seq). Results confirmed strong upregulation of many transposon families upon Su(var)2-10 knockdown (Fig 2B). Consistent with its stronger knockdown efficiency and phenotypic effect, the Sv210-2 hairpin induced broader and stronger TE upregulation (Fig. 2B, S1A). We validated these results by performing a further RNA-seq experiment of shSv210-2 and control ovaries in two independent biological replicates. Results showed a highly reproducible upregulation of TEs (Fig. S1B). Differential expression analysis showed that 60% (116 of 195) of all TE families were more than 2-fold upregulated, with 30% more than 4-fold upregulated (FDR<0.05; Fig. S1C). These data therefore suggest that Su(var)2-10 plays a major role in supressing the activity of many different transposon families in the *Drosophila* germline. As the piRNA pathway plays a central role in TE silencing in the germline(LeThomas et al., 2013; Rozhkov et al., 2013), we compared transposon expression upon Su(var)2-10 and Piwi GLKD. This analysis revealed that Su(var)2-10 and Piwi have similar target repertoires (Fig. 2C), suggesting that Su(var)2-10 may be a novel player in piRNA-mediated transposon repression in the germline. In addition, Su(var)2-10 GLKD affected the expression of approximately 10% of the host genes (Fig S1C). Detailed investigation of the role of Su(var)2-10 in host transcriptome regulation revealed complex TE-dependent and TE-independent effects that we describe in detail in the accompanying manuscript (Ninova et al., accompanying manuscript).

**Figure 2.**
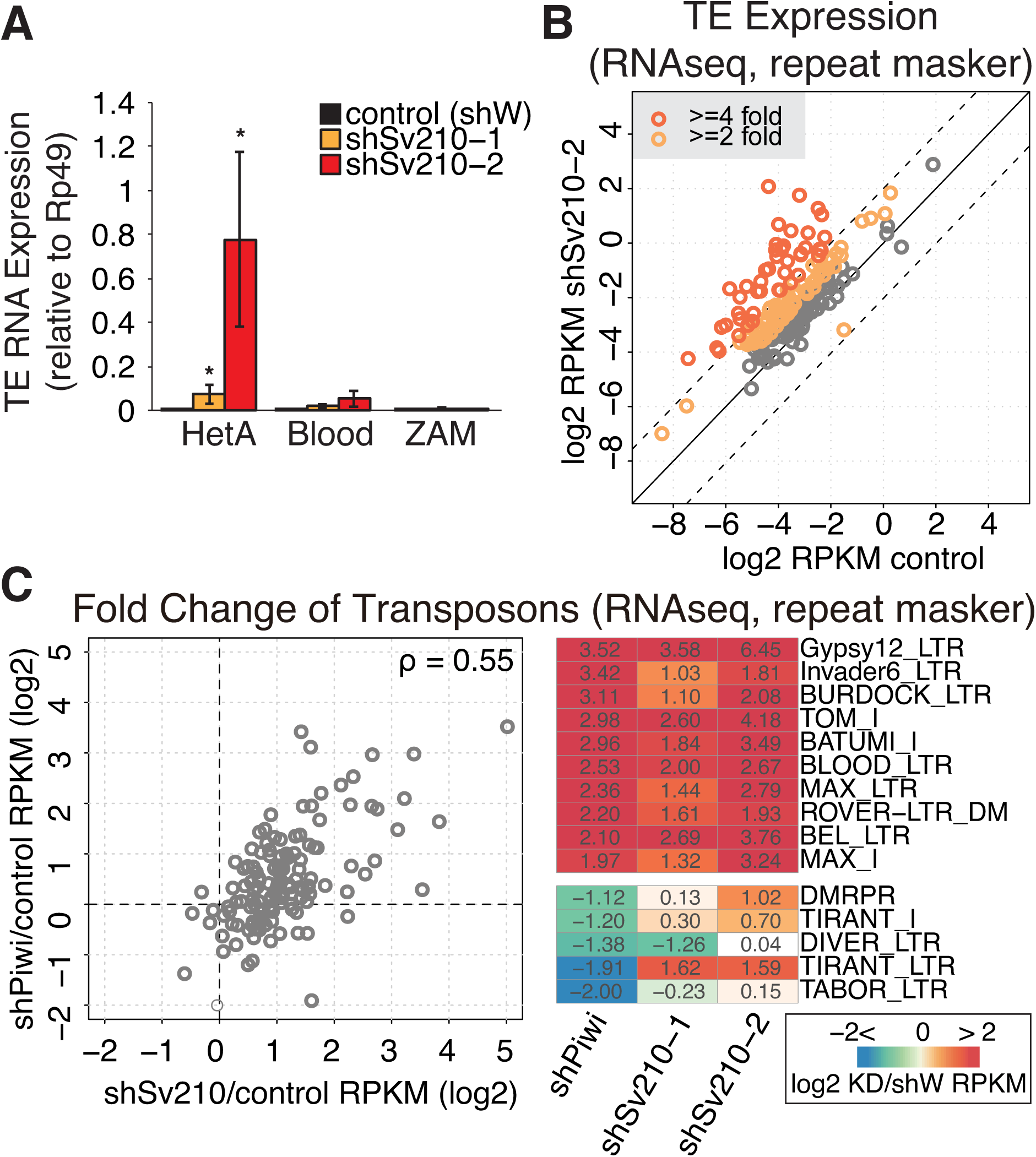
Germline depletion of Su(var)2-10 leads to transcriptional upregulation of transposons similar to those in Piwi depletion. (A)Depletion of Su(var)2-10 in germline cells de-represses germline-specific transposons. Relative HetA (germline specific), Blood (intermediate) and ZAM (soma specific) TE levels in control and Su(var)2-10 GLKD ovaries as estimated by RT-qPCR. *Rp49* mRNA was used as an endogenous control; error bars show the standard deviation from three biological replicas. Statistical significance was determined by two-tailed Student’s t-test. Significant increase compared to the control is marked *p<0.05. (B)Su(var)2-10 depletion leads to global transposon derepression. Scatter plot shows log2-tansformed RPKM values for TEs (RepeatMasker) in RNA-seq data from ovaries expressing the shSv210-2 hairpin (Y-axis) versus the shW control hairpin (X-axis), under the control of the MT-Gal4 driver. Dashed lines indicate 4-fold change. (C)Germline-specific Piwi and Su(var)2-10 knockdowns lead to de-repression of similar TEs. (Left) Scatter plot shows the fold changes in TE expression upon Piwi (Y-axis) and Su(var)2-10 (X-axis) knockdowns estimated by RNA-seq. Data points reflect log2-transformed ratios of TE RPKM values from knockdown versus control ovaries (shPiwi/shW and shSv210/shW). The Spearman’s correlation coefficient (rho) is shown. (Right) Heatmap shows log2-transformed fold changes of RNA expression in knockdown versus control ovaries for the 10 most upregulated and 5 most downregulated TEs in Piwi-depleted ovaries.

### Su(var)2-10 depletion correlates with loss of repressive histone marks and gain of active marks at transposon sequences

In the nucleus, piRNA-guided PIWI proteins induce co-transcriptional repression associated with trimethylation of histone H3 lysine 9 (H3K9me3), a repressive histone modification that serves as a binding site for heterochromatin protein 1 (HP1)(Bannister et al., 2001; Jacobs et al., 2001; Lachner et al., 2001). To test whether Su(var)2-10 is involved in transcriptional silencing of transposons, we analysed the effect of Su(var)2-10 depletion on the levels of H3K9me3 and HP1a by ChIP-seq.

In wild-type flies we found the high H3K9me3 levels in heterochromatic genomic regions, including pericentromeric and telomeric regions of the chromosomal arms (Fig. 3A, black bars). In addition, we identified H3K9me3 peaks scattered across euchromatic regions. Regions with elevated H3K9me3 signal are highly enriched in transposon sequences: a median of 70% of their sequence annotated by RepeatMasker (Fig. S2), while transposons occupy less than 20 % of whole reference genome sequence(Kaminker et al., 2002). A subset of the detected H3K9me3 peaks in euchromatin lack transposon sequences in the reference genome within 10 kb proximity. However, transposition of mobile elements might lead to new transposon insertions that are absent in the reference genome sequence, but might be present in the genome of the strain used in our experiments. Indeed, analysis of *de novo* TE insertions in the genome of strains used in this study using the TIDAL pipeline(Rahman et al., 2015) identified 119 novel transposon integrations, which were absent in the reference genome, residing within 5 kb of euchromatic H3K9me3 peaks (n=479). The association of H3K9me3 islands with non-reference TE integration is more frequent than expected by chance (p<1×10^−6^, permutation test). Thus, the H3K9me3 mark correlates with TE sequences in both eu- and heterochromatin.

**Figure 3.**
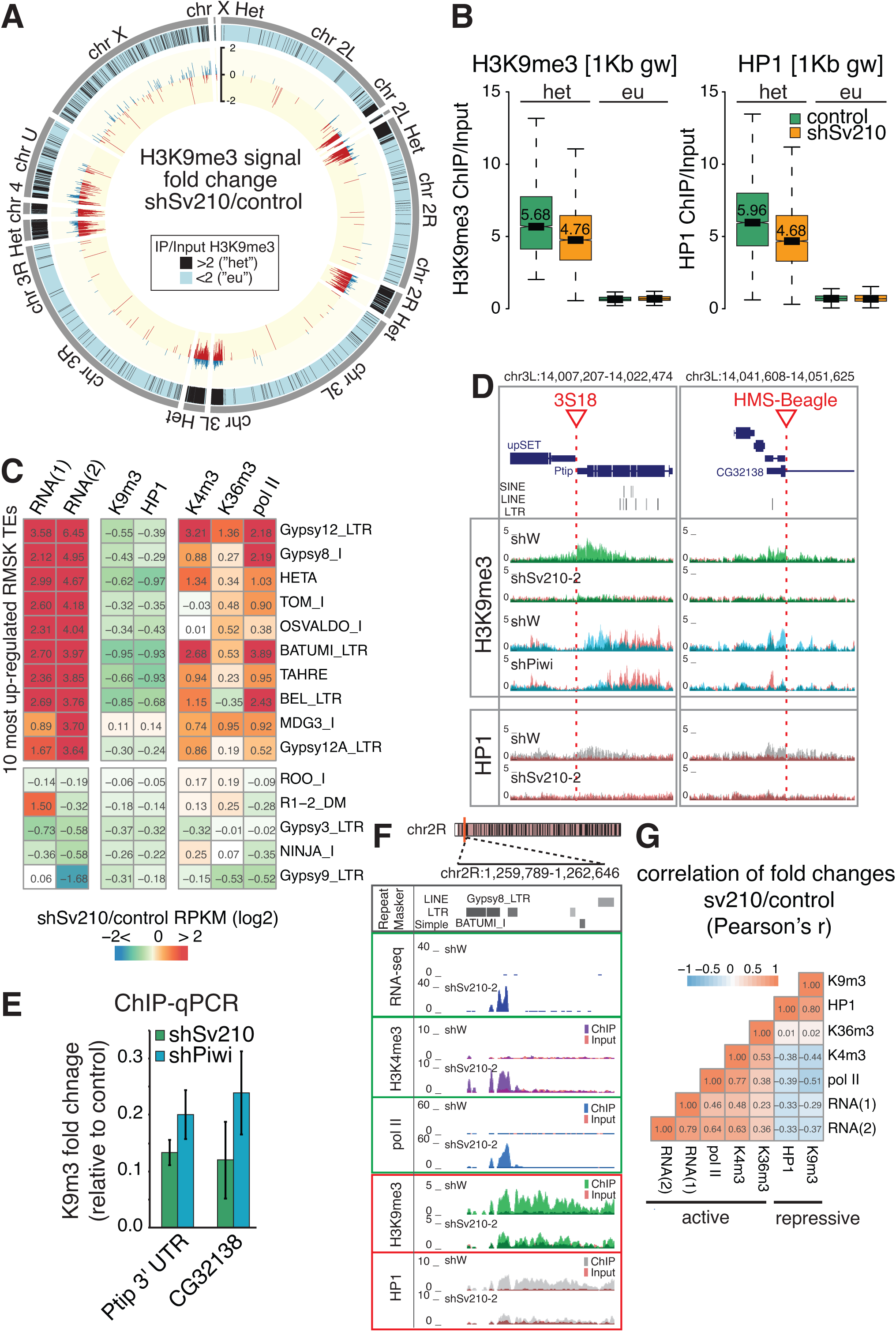
Su(var)2-10 depletion induces ubiquitous H3K9me3 and HP1a loss associated with transposon de-repression. (A)Genome-wide distribution and fold change of the H3K9me3 mark upon Su(var)2-10 depletion. Outer grey tiles represent different chromosome arms. H3K9me3 signal is calculated for 1 Kb genomic windows as log2-normalized ratio of ChIP-seq to Input RPKM values. Data is average from two biological replicates from control (shW) ovaries. Regions with H3K9me3 ChIP to Input signal >2 are defined as heterochromatic (het) and marked by black tiles. Inner histogram shows the log2-transformed fold change of H3K9me3 signal in Su(var)2-10 KD versus control (shW) ovaries in ‘het’ regions. Positive and negative fold changes are color-coded in blue and red, respectively. Values for euchromatic intervals are not displayed. Circle plot is generated with Circos(Krzywinski et al., 2009), using the dm3 assembly of the *D. melanogaster*genome. (B)Su(var)2-10 depletion induces loss of H3K9me3 and HP1a from heterochromatic regions. Boxplots show the distributions of H3K9me3 (left) and HP1a (right) signal in H3K9me3-enriched (het, >2fold H3K9me3 enrichment in control ovaries), and euchromatic 1Kb genomic windows for Su(var)2-10 GLKD and control (shW) ovaries. Data is average from two biological replicates. Median values are shown. (C)TE upregulation in Su(var)2-10 GLKD correlates with loss of repressive chromatin marks and gain of active transcription marks. Heatmap shows log2-transformed fold changes between Su(var)2-10 GLKD and control (shW) ovaries in RNA-seq and ChIP-seq datasets as indicated. Values for the 10 most upregulated and 5 most downregulated TEs on RNA level upon Su(var)2-10 depletion are shown. (D)H3K9me3 and HP1a loss upon Su(var)2-10 and Piwi knockdown at genomic regions adjacent to novel TE insertions. UCSC browser tracks display three examples of euchromatic loci adjacent to novel non-reference TE insertions. Top diagrams show genomic coordinates, RefSeq annotations and RepeatMasker regions. Red arrows indicate the position of the non-reference TE detected by TIDAL(Rahman et al., 2015). Histograms show RPM-normalized ChIP and Input signal for control (shW) and Su(var)2-10 knockdown ovaries (shSv210-2), and control (shW) and Piwi knockdown ovaries (shPiwi). Only uniquely mapping reads are shown. For each ChIP experiment, IP and input signal tracks are overlaid. (E)Loss of H3K9me3 is similar upon Su(var)2-10 and Piwi GLKD. Changes in H3K9me3 mark upon Su(var)2-10 and Piwi germline knockdowns for three genomic loci adjacent to the non-reference TE insertions shown in (D) were assessed by ChIP-qPCR. ChIP signal is normalized to input, and the ratio between knockdown and control (shW) ovaries is shown. Error bars show the standard deviation of two biological replicates. (F)Su(var)2-10 GLKD causes increase of active transcription marks and loss of repressive transcription marks proximal to annotated TEs. UCSC browser tracks display and example of changes in transcription and chromatin marks near TE repeats upon Su(var)2-10 depletion. Histograms show RPM-normalized RNA-seq and ChIP-seq tracks for control (shW) and shSv210-2 knockdown ovaries. Only uniquely mapping reads are shown. For each ChIP experiment, IP and input signal tracks are overlaid. (G)Correlation between changes in active and repressive marks upon Su(var)2-10 GLKD. Heatmap shows all-versus-all correlation coefficients for fold changes upon Su(var)2-10 knockdown of steady state RNA levels (RNA-seq of GLKD with two different hairpins against shSu(var)2-10), active transcription marks (ChIP-seq), and repressive marks (ChIP-seq) at TE families annotated by RepeatMasker. ChIP-seq fold change values are calculated as average of two biological replicates.

Su(var)2-10 depletion caused a genome-wide reduction of the H3K9me3 mark: the majority (∼80%) of H3K9me3-enriched genomic intervals displayed a decreased H3K9me3 signal upon Su(var)2-10 GLKD (Fig. 3A,B). Consistent with loss of H3K9me3, depletion of Su(var)2-10 also induced a decrease in HP1a level in these genomic regions (Fig. 3B, right). In line with the global loss of H3K9me3, analysis revealed a widespread decrease of H3K9me3 and HP1a signal on individual TE families, especially at transposons that show a strong derepression upon Su(var)2-10 knockdown (Fig. 3C). Regions flanking non-reference TE insertions in euchromatin also show prominent loss of H3K9me3 and associated HP1a upon Su(var)2-10 GLKD (Fig. 3D). Notably, the same regions exhibited a reduction of H3K9me3 upon Piwi depletion, as demonstrated by both ChIP-seq and qPCR (Fig. 3D,E), indicating that they are controlled by both Piwi and Su(var)2-10.

We also assessed the effect of Su(var)2-10 GLKD on chromatin marks associated with active transcription, including the H3K4me3 mark that is present at active promoters, the elongation mark H3K36me3, as well as RNA polymerase II occupancy. Genome-wide ChIP-seq analysis of two biological replicas revealed that depletion of Su(var)2-10 results in an increase of active transcription marks over transposon sequences and flanking regions (Fig. 3C,F). These changes correlate with the increase of transposon RNA levels, and the decrease of H3K9me3 and HP1a (Fig. 3G). Together, the loss of repressive histone marks and the gain of active marks upon Su(var)2-10 GLKD imply that Su(var)2-10 controls TE expression in the germline through transcriptional silencing.

### Su(var)2-10 interacts with the piRNA/Piwi/Panx silencing complex and is required for its ability to induce transcriptional repression

The molecular mechanism of Piwi-induced transcriptional silencing remains poorly understood. Recent studies identified two proteins, Arx and Panx, that form a complex with Piwi and are required for transcriptional silencing of Piwi targets(Donertas et al., 2013; Muerdter et al., 2013; Ohtani et al., 2013; Sienski et al., 2015; Yu et al., 2015). Recruitment of Panx to a reporter locus in a piRNA-independent manner results in transcriptional repression and H3K9me3 deposition at the reporter(Sienski et al., 2015; Yu et al., 2015), providing a model system to study Piwi-induced silencing. We took advantage of this system to test whether Su(var)2-10 is required for Piwi repression downstream of Panx. In agreement with previous reports(Sienski et al., 2015; Yu et al., 2015), Panx tethering resulted in local deposition of the H3K9me3 mark at the reporter locus (Fig 4A). Knockdown of Su(var)2-10 reduced the ability of Panx to induce H3K9 trimethylation (Fig 4A), indicating that Su(var)2-10 is required for Panx-induced silencing. Thus, Su(var)2-10 acts downstream of Panx and is required for Panx-induced H3K9me3 deposition.

**Figure 4.**
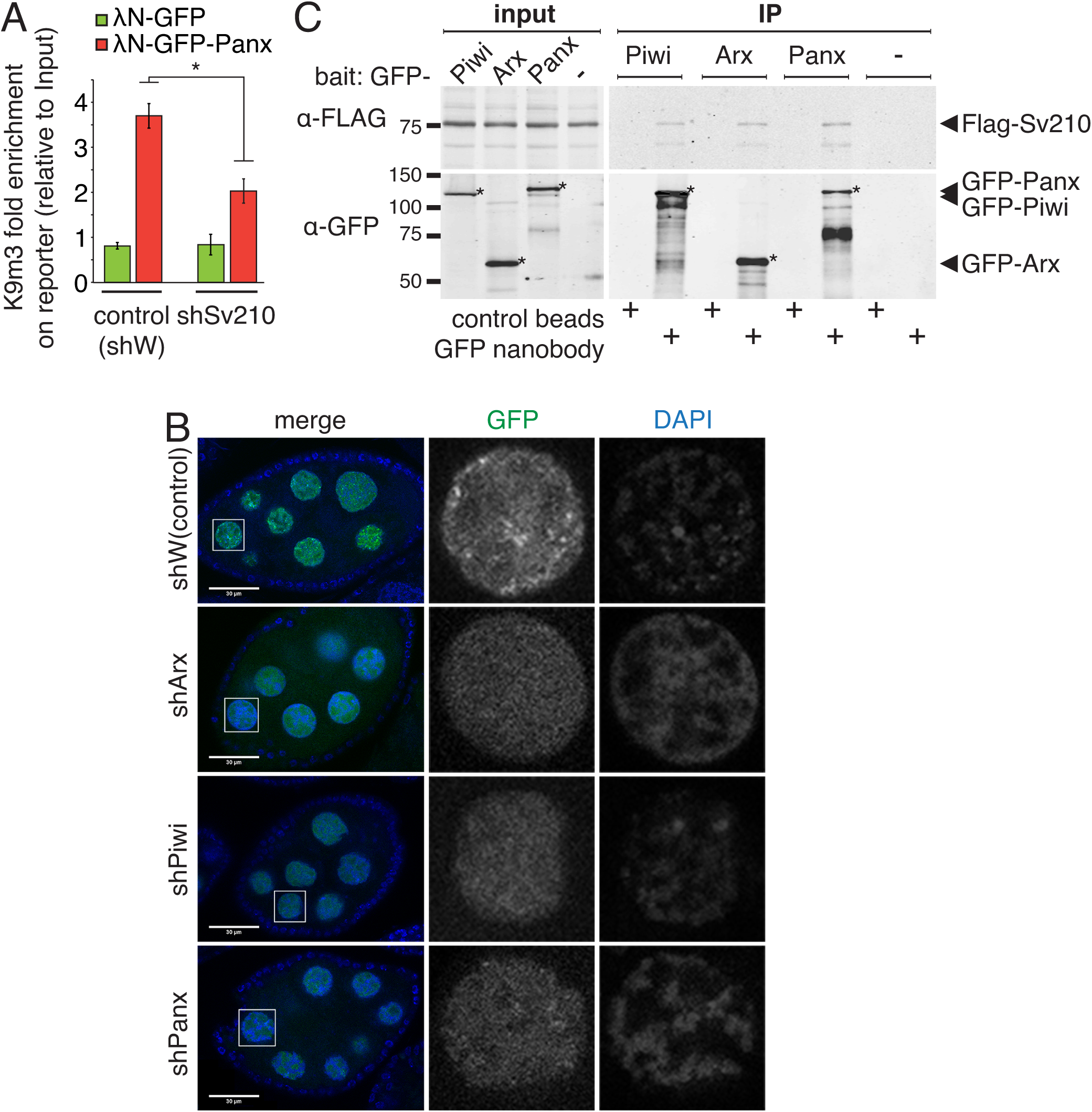
Su(var)2-10 genetically and biochemically interacts with Piwi, Arx and Panx. (A)Su(var)2-10 knockdown abolishes H3K9me3 deposition induced by Panx tethering to luciferase reporter(Yu et al., 2015) locus. Bar plots show H3K9me3 levels (ChIP-qPCR) at reporter locus upon tethering of Panx or GFP to reporter in control or Su(var)2-10 knockdown ovaries. Data is normalized to Rp49 as endogenous control, and fold enrichment is calculated as the ratio of signal in ChIP to signal in input. Error bars show the standard deviation from two biological replicates. Statistical significance is estimated by two-tailed Student’s t-test; *p<0.05. (B)Nuclear localization of Su(var)2-10 depends on Arx, Panx and Piwi. Images show nurse cell nuclei of ovaries from flies expressing GFP-Su(var)2-10 (green channel) and short hairpins against Asterix (shArx), Piwi (shPiwi), Panoramix (shPanx) or *white* (shW, control) under the control of MT-gal4 driver. Nuclei are stained with DAPI (blue channel). Scale bar 30 μm. (C)Su(var)2-10 interacts with Piwi, Arx and Panx. Protein lysates from ovaries overexpressing MT-gal4 driven FLAG-Su(var)2-10 and GFP-fusion Piwi, Arx or Panx, were used for co-immunoprecipitation using GFP nanotrap or control beads. Samples expressing FLAG-Su(var)2-10 only but no GFP partner (no bait) were used as an additional control. Western blots were probed with anti-FLAG antibody to detect Su(var)2-10 and anti-GFP antibody to detect bait proteins.

To further explore the genetic interactions between Su(var)2-10 and components of the Piwi-induced transcriptional silencing complex, we analysed the subcellular localization of Su(var)2-10 in wild-type flies and upon disruption of Piwi-induced transcriptional silencing. Consistent with a previous report that Su(var)2-10 associates with chromatin(Hari et al., 2001), we found that GFP-tagged Su(var)2-10 localizes to the nucleus of nurse cells, where it concentrates at discrete foci, possibly indicating binding at specific genomic sites. Depletion of Piwi, Panx or Arx in germ cells altered the localization of GFP-Su(var)2-10 to a uniform distribution in the nucleus (Fig 4B). The requirement of Piwi and its auxiliary factors Arx and Panx for proper localization of Su(var)2-10 further suggests that Su(var)2-10 acts downstream of these factors.

To test if Su(var)2-10 physically interacts with components of the Piwi/Arx/Panx complex, we employed a co-immunoprecipitation assay using ovaries from transgenic flies expressing tagged proteins. Su(var)2-10 co-purified with all three factors, Piwi, Panx and Arx (Fig. 4C), indicating that Su(var)2-10 associates with the Piwi/piRNA transcriptional silencing complex. The interaction of Su(var)2-10 with Piwi, Panx and Arx was further validated by co-expression and co-immunoprecipitation of tagged proteins from *Drosophila* S2 cell lines (Fig. S3A-C). Overall, our results indicate that Su(var)2-10 interacts with the Piwi-induced silencing complex both genetically and physically, and that it is required for Piwi-induced transcriptional repression.

### Recruitment of Su(var)2-10 to a genomic locus induces transcriptional repression and accumulation of the H3K9me3 mark

The requirement of Su(var)2-10 for transposon repression and Panx-induced reporter silencing suggest that Su(var)2-10 plays a role in transcriptional repression downstream of target recognition by the piRNA/Piwi/Panx complex. To test if Su(var)2-10 is able to induce local transcriptional repression when recruited to chromatin, we tethered Su(var)2-10 to a reporter locus in a piRNA-independent manner by fusing it to the λN RNA-binding domain, which has a high affinity for BoxB hairpins encoded in the 3’UTR of a reporter RNA (Fig. 5A). Tethering of λN-GFP-Su(var)2-10 caused severe (∼130 fold) decrease in reporter mRNA expression, indicating that recruitment of Su(var)2-10 is sufficient to induce strong repression (Fig. 5B). Recruitment of Su(var)2-10 also resulted in an increase in the repressive H3K9me3 mark and a decrease in Pol II occupancy on the reporter as measured by ChIP-qPCR (Fig. 5C, D). Similar results were obtained when Su(var)2-10 was recruited to an alternate reporter containing a different sequence and integrated at another locus, indicating that the results are independent of reporter sequence and local genomic environment (Fig. S4A). These results indicate that recruitment of Su(var)2-10 to a genomic locus induces strong transcriptional repression associated with accumulation of the repressive H3K9me3 mark.

**Figure 5.**
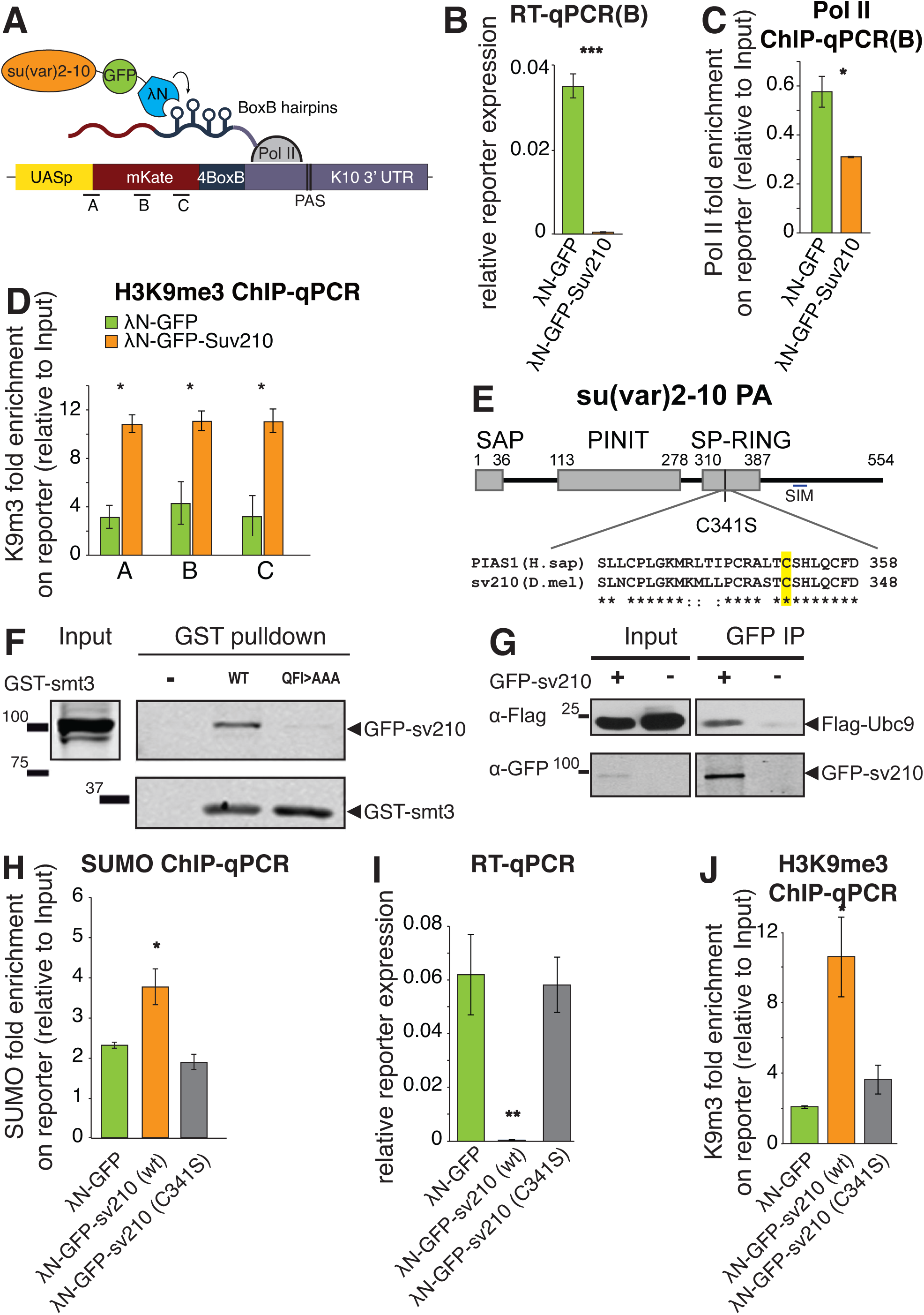
Su(var)2-10 recruitment induces transcriptional repression that depends on the SUMO pathway. (A)Schematic diagram of the reporter used to study the effect of Su(var)2-10 recruitment to RNA. GFP-Su(var)2-10 or just GFP (control) are fused to the RNA-biding AN domain, which has high affinity for BoxB hairpins and the mKate reporter encoding 4 BoxB hairpins in the 3’UTR are co-expressed in ovaries resulting in Su(var)2-10 recruitment to the reporter’s nascent transcript. Regions A-C denote the positions of RT-qPCR amplicons. (B)Tethering of Su(var)2-10 leads to reduced reporter mRNA level. Boxplot shows reporter expression levels (region B) in ovaries expressing AN-GFP-Su(var)2-10 or AN-GFP control as estimated by RT-qPCR. Error bars show the standard deviation from 3 biological replicates. Statistical significance is estimated by two-tailed Student’s t-test. ***p<0.001 (C)Tethering of Su(var)2-10 leads to decreased Pol II occupancy at the reporter locus. Boxplot shows Pol II occupancy at the reporter locus in ovaries expressing AN-GFP-Su(var)2-10 or AN-GFP control as estimated by ChIP-qPCR. Data is normalized to Rp49 as endogenous control, and fold enrichment is calculated as the ratio of signal in ChIP to signal in input. Error bars show the standard deviation from two biological replicates. Statistical significance is estimated by two-tailed Student’s t-test. *p<0.05 (D)Tethering of Su(var)2-10 leads to H3K9me3 deposition at reporter locus. Boxplot shows H3K9me3 levels at the reporter locus in ovaries expressing AN-GFP-Su(var)2-10 or AN-GFP control as estimated by ChIP-qPCR (for regions A-C). Data is normalized to Rp49 as endogenous control, and fold enrichment is calculated as the ratio of signal in ChIP to signal in input. Error bars show the standard deviation from two biological replicates. Statistical significance is estimated by two-tailed Student’s t-test. *p<0.05 (E)Schematic diagram of *Drosophila* Su(var)2-10 protein structure (PA isoform). Grey boxes mark conserved domains. Local alignment of the SP-RING domain between human PIAS1 and Su(var)2-10 is shown, highlighting the catalytic cysteine residue identified in human PIAS1(Kahyo et al., 2001). Sequences were extracted from UniProt and aligned by MUSCLE (Edgar, 2004). (F)Su(var)2-10 interacts with SUMO. S2 cell lysates expressing GFP-Su(var)2-10 were incubated with bacterially expressed GST-SUMO (wild type), interaction-deficient mutant GST-SUMO (QFI>AAA), or no bait control. GST-SUMO was affinity purified using glutathione sepharose beads and GFP-Su(var)2-10 was detected by Western blotting using an anti-GFP antibody. (G)Su(var)2-10 interacts with Ubc9. S2 cell lysates expressing FLAG-tagged Ubc9 and GFP-Su(var)2-10 were analysed by immunoprecipitation using GFP nanotrap beads. FLAG-Ubc9 co-purifies with GFP-Su(var)2-10. (H)Su(var)2-10 tethering promotes SUMO enrichment at the reporter locus. Boxplot shows SUMO levels at reporter locus in ovaries expressing AN-GFP-Su(var)2-10 (wild type or mutant) or AN-GFP control as estimated by ChIP-qPCR (region A). Wild type but not C341S Su(var)2-10 mutant induces SUMO enrichment. Data is normalized to Rp49 as endogenous control, and fold enrichment is calculated as the ratio of signal in ChIP to signal in input. Error bars show the standard deviation from two biological replicates. Statistical significance is estimated by one-way ANOVA followed by Tukey’s test; *p<0.05. (I)Su(var)2-10 mutant (C341S) is unable to repress reporter transcription. AN-GFP control, AN-GFP-Su(var)2-10 (wild type) or AN-GFP-Su(var)2-10 C341S mutant were co-expressed in ovaries with mKate-4BoxB reporter. Bar plots show reporter expression levels (region B) measured by RT-qPCR. Tethering of wild type, but not catalytically inactive AN-GFP-Su(var)2-10 induces reporter repression. Error bars show the standard deviation of three biological replicates. Statistical significance is estimated by one-way ANOVA followed by Tukey’s test; **p<0.01. (J) Catalytically inactive Su(var)2-10 mutant (C341S) is unable to induce H3K9me3 deposition. Bar plots show H3K9me3 levels at the mKate-4BoxB reporter upon tethering of AN-GFP control, AN-GFP-Su(var)2-10 (wild type) or AN-GFP-Su(var)2-10 C341S as measured by ChIP-qPCR (region B). Recruitment of wild type, but not catalytically inactive AN-GFP-Su(var)2-10 induces H3K9me3 enrichment. ChIP data is analysed as in D. Error bars show standard deviation of two biological replicates. Statistical significance is estimated by one-way ANOVA followed by Tukey’s test; *p<0.05.

### Su(var)2-10 is a SUMO E3 ligase

Su(var)2-10 is a member of the PIAS (Protein Inhibitor of Activated STAT) protein family(Betz et al., 2001; Hari et al., 2001; Mohr and Boswell, 1999). PIAS proteins were first identified as inhibitors of the JAK-STAT pathway(Chung et al., 1997), however, subsequent genetic and biochemical studies showed that members of this family in yeast and mammals function in the SUMO pathway, which covalently attaches SUMO (Small Ubiquitin-like Modifier) to proteins to modify their activity(Johnson and Gupta, 2001; Kahyo et al., 2001; Kotaja et al., 2002; Sachdev et al., 2001; Schmidt and Muller, 2002; Takahashi et al., 2001). Yeast and mammalian PIAS proteins interact with SUMO and the E2 SUMO-conjugating enzyme Ubc9, and facilitate the transfer of SUMO from Ubc9 to substrates, thereby acting as SUMO E3 ligases(Johnson and Gupta, 2001; Kahyo et al., 2001; Kotaja et al., 2002; Sachdev et al., 2001; Schmidt and Muller, 2002; Takahashi et al., 2001). *Drosophila* Su(var)2-10 emerged as a factor involved in the SUMO pathway in a genetic screen(Stielow et al., 2008a), however, its biochemical functions were not well characterized. Thus, we decided to ask whether Su(var)2-10 has a SUMO E3 ligase activity, and whether this activity is required for transcriptional repression.

*D. melanogaster* Su(var)2-10 has all the conserved domains present in yeast and mammalian members of the PIAS family, including the Siz/PIAS RING (SP-RING) domain responsible for interaction with Ubc9 and the E3 ligase activity in mammals(Kahyo et al., 2001), as well as a predicted SUMO interaction motif (SIM) at its C-terminus (Fig. 5E). First, we tested if Su(var)2-10 interacts with SUMO. Su(var)2-10 binds purified SUMO protein *in vitro* (Fig. 5F, S4B). In contrast, Su(var)2-10 does not interact with a SUMO mutant generated by changing three conserved residues (Q26A, F27A, I29A) essential for binding to the SUMO interaction motif(Zhu et al., 2008), confirming specificity of the interaction (Fig. 5F). Next, we tested the interaction between Su(var)2-10 and the E2 SUMO ligase Ubc9. Co-immunoprecipitation of tagged proteins expressed in S2 cells revealed that Su(var)2-10 binds Ubc9 (Fig. 5G). The interactions with SUMO and Ubc9 indicate that Su(var)2-10 indeed acts in the SUMO pathway and likely has a conserved function as a SUMO E3 ligase.

PIAS proteins and other SUMO E3 ligases were found to be able to self-SUMOylate (Ivanov et al., 2007; Kotaja et al., 2002; Schmidt and Muller, 2002). For example, auto-SUMOylation of the KAP1 SUMO E3 ligase was shown to be an important regulatory step for silencing complex recruitment in mammalian systems(Ivanov et al., 2007). To test if Su(var)2-10 can be SUMOylated, we employed an *in vitro* assay using purified Su(var)2-10, SUMO, E1 and E2 enzymes. In the presence of ATP, we observed a higher molecular weight Su(var)2-10 band suggesting that Su(var)2-10 is itself a substrate of SUMO modification (Fig. S4C).

### The repressive function of Su(var)2-10 depends on the SUMO pathway

To explore the ability of Su(var)2-10 to promote SUMOylation on chromatin *in vivo*, we assessed the level of SUMO at the reporter locus upon Su(var)2-10 tethering. ChIP-qPCR using an antibody against the endogenous *Drosophila* SUMO protein (smt3)(Gonzalez et al., 2014) showed that tethering of Su(var)2-10 led to an increased SUMO signal at the reporter locus, strongly supporting its function as a SUMO E3 ligase that modifies chromatin targets (Fig. 5H).

Next, we asked if the SUMO E3 ligase activity of Su(var)2-10 is important for its function in transcriptional repression. Studies of human PIAS1 revealed that its interaction with Ubc9 and its SUMO E3 ligase activity depend on the SP-RING domain, and a point mutation of a single cysteine residue in this domain abolishes this activity(Kahyo et al., 2001; Munarriz et al., 2004). The SP-RING domain is well conserved between *Drosophila* Su(var)2-10 and human PIAS1, including the aforementioned cysteine residue (Fig. 5E). Thus, we generated a C341S point mutant of Su(var)2-10 to study the role of the SUMO E3 ligase activity in transcriptional repression. Tethering of Su(var)2-10 C341S mutant failed to promote SUMOylation at the reporter locus, confirming that this mutation abolishes its E3 ligase function (Fig. 5H). We next asked whether the Su(var)2-10 C341S mutant is able to induce reporter repression and H3K9me3 deposition. Tethering of wild-type Su(var)2-10 reproducibly induced over 100-fold reporter repression. In contrast, tethering of the Su(var)2-10 C341S did not affect reporter expression or H3K9me3 level at its genomic locus, indicating that the SUMO-ligase activity of Su(var)2-10 is essential for its function in transcriptional repression (Fig. 5I,J). Next, we addressed the role of different Su(var)2-10 domains in its repressive activity (Fig. S4D). As expected, complete deletion of the SP-RING domain impaired reporter silencing. Deletion of the PINIT domain also abolished the silencing activity completely. Conversely, deletion of the SAP domain, which was proposed to bring PIAS proteins to some of its targets through binding to DNA(Reindle et al., 2006), caused reporter repression at levels comparable of those induced by the wild-type protein. Overall, these results indicate that the ability of Su(var)2-10 to induce transcriptional repression depends on its SUMO E3 ligase activity.

To directly test if SUMO is required for Su(var)2-10-induced transcriptional repression, we studied the ability of wild-type Su(var)2-10 to induce silencing upon knockdown of the single SUMO gene *(smt3)* encoded in the *Drosophila* genome. Depletion of SUMO in germ cells released Su(var)2-10-induced reporter repression ∼10 fold (Fig. 6A). Thus, transcriptional silencing caused by recruitment of Su(var)2-10 to chromatin is dependent on the SUMO pathway.

**Figure 6.**
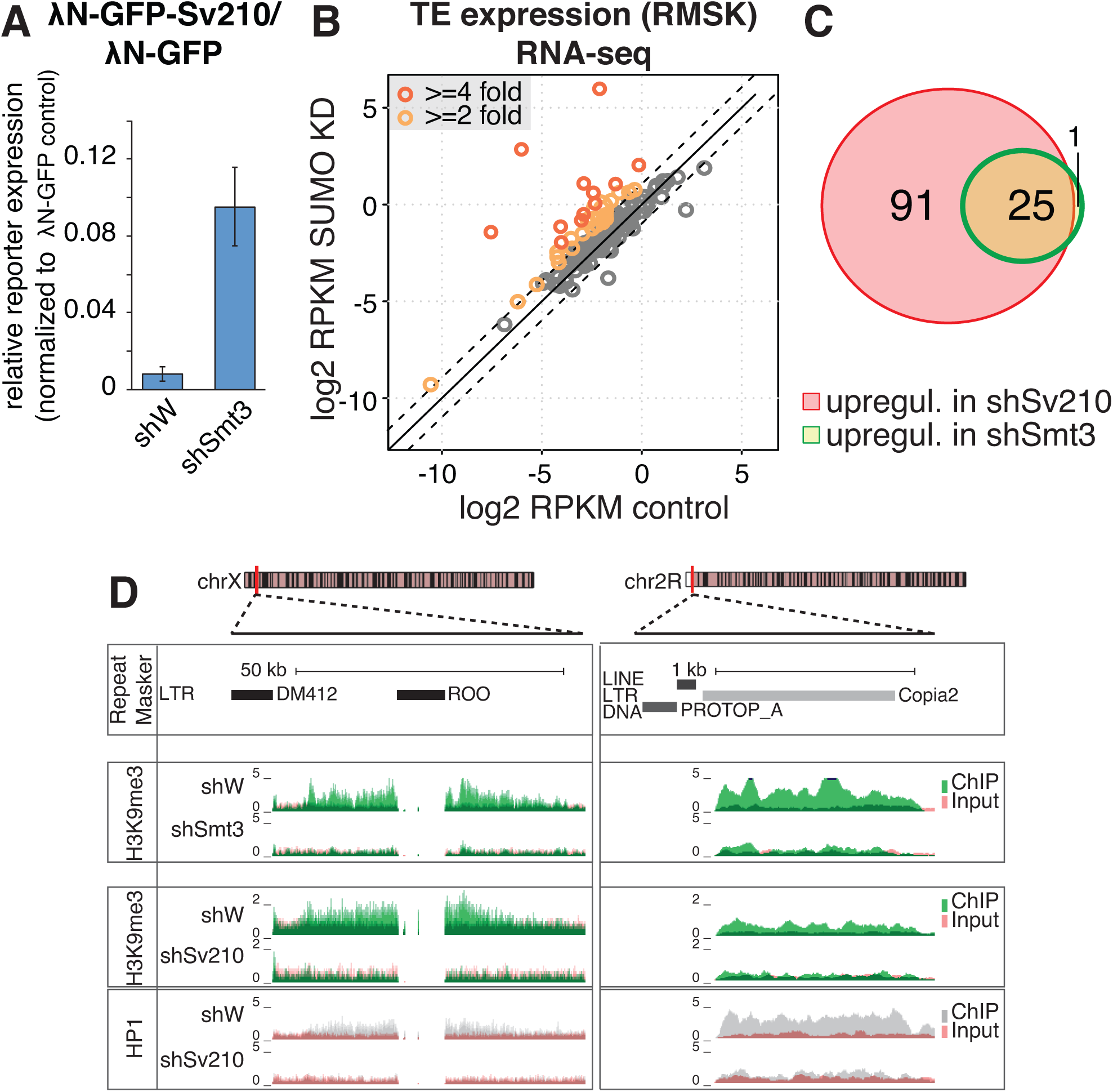
SUMO is required for Su(var)2-10 mediated reporter silencing and global regulation of TEs. (A)SUMO is required for Su(var)2-10 mediated reporter silencing. AN-GFP or AN-GFP-Su(var)2-10 were recruited to the mKate-4BoxB reporter (Fig 5A) in SUMO knockdown (shSmt3) or control (shW) ovaries. Reporter expression is measured by RT-qPCR and normalized to rp49 as endogenous control. Bar plots show the ratio of reporter expression in AN-GFP-Su(var)2-10 tethering to AN-GFP tethering for each GLKD. Error bars show the propagated standard deviations from three biological replicates for each condition. (B)SUMO depletion leads to global transposon de-repression. Scatter plots show log2-tansformed RPKM values for TEs (RepeatMasker annotations) in RNA-seq data from ovaries from shSmt3 (SUMO knockdown) (Y-axis) versus shW control lines (X-axis). Dashed lines indicate 2-fold change. Data is average from two biological replicates. (C)TE targeted by SUMO and Su(var)2-10 overlap. Venn diagram shows the numbers and the overlap of significantly upregulated transposon families between control and GLKD of SUMO and Su(var)2-10 ovaries as determined by DESeq2 (FDR<0.05, log2 fold change >=2). Nearly all TEs affected by SUMO knockdown are also affected by Su(var)2-10 knockdown in the germline. (D)SUMO depletion leads to loss of H3K9me3 signal at transposon loci. UCSC browser tracks display examples of H3K9me3 and HP1a loss upon Su(var)2-10 and SUMO knockdown at genomic regions enriched in TE elements. Top diagrams show genomic position and RepeatMasker annotated TEs. Histograms show RPM-normalized ChIP and Input signal for control (shW), Su(var)2-10 GLKD and SUMO (shSmt3) GLKD ovaries for indicated histone marks. Only uniquely mapping reads are shown. For each ChIP experiment, IP and input signal tracks are overlaid.

To determine whether the SUMO pathway is required for piRNA/Piwi mediated transcriptional silencing, we investigated the effect of SUMO depletion on expression of transposons in the germline. Depletion of *smt3* in germ cells resulted in sterility, phenocopying the effect of knockdowns of *Su(var)2-10* and other piRNA pathway mutants. Global RNA-seq followed by DESeq2 analysis revealed upregulation of 26 transposable element families (>2-fold increase, FDR<0.05) (Fig. 6B). Notably, the set of transposons that are depressed upon SUMO depletion nearly completely overlaps with elements upregulated in *Su(var)2-10* knockdown, suggesting that Su(var)2-10 and SUMO act on the same targets (Fig. 6C). Furthermore, global ChIP-seq analysis showed loss of the H3K9me3 mark from transposon sequences and flanking regions upon *smt3* GLKD, as exemplified in Fig. 6D. Thus, SUMO is crucial for transcriptional repression of TEs in the germline.

### Transcriptional repression induced by Su(var)2-10 requires the histone methyltransferase complex SetDB1/Wde

H3K9me3 deposition induced by Piwi and Panx requires the activity of the methyltransferase SetDB1 also known as Eggless(Sienski et al., 2015; Yu et al., 2015). We found that transcriptional repression by Su(var)2-10 is dependent on the SUMO pathway. Previous reports indicate that the mammalian homolog of SetDB1, as well as its co-factor MCAF1/ATF7IP, have SUMO-interaction motifs (SIM) (Ivanov et al., 2007; Stielow et al., 2008b; Thompson et al., 2015; Uchimura et al., 2006). To explore whether the SUMO pathway plays a role in recruitment of SetDB1 to chromatin in *Drosophila*, we analysed the sequences of fly SetDB1 and its conserved co-factor called Windei (Wde) (Koch et al., 2009). Computational analysis(Zhao et al., 2014) identified several canonical SIMs in both proteins (Fig. 7A,S5A). Furthermore, SIMs in SetDB1 and Wde are conserved between *D. melanogaster* and other Drosophilid species separated by 30-60 My of evolution (Fig. 7A,S5A). While the fly and human proteins have little sequence homology outside of conserved domains, the relative position of some SIMs is preserved (Fig. 7A,S5A). To test if *Drosophila* SetDB1 and Wde interact with SUMOylated proteins, we co-transfected S2 cells with tagged SetDB1 or Wde and SUMO followed by purification of SetDB1 and Wde complexes. Western blotting showed that both SetDB1 and Wde co-purify with several SUMOylated proteins (Fig. 7B). Thus, Drosophila SetDB1 and Wde possess conserved SUMO-interaction motifs and interact with SUMOylated proteins. To further explore the interaction between Wde and SUMO, we expressed SIM-containing fragments of Wde in S2 cells and incubated the lysate with purified GST-SUMO. Wde fragments co-purified with wild type, but not triple point mutant (Q26A, F27A, I29A) SUMO. Mutations in the SIM of Wde abolished its interaction with wild-type SUMO, indicating that Wde binds SUMO directly via its SIM (Fig. 7C). Thus, the SetDB1/Wde complex directly and specifically binds SUMO.

**Figure 7.**
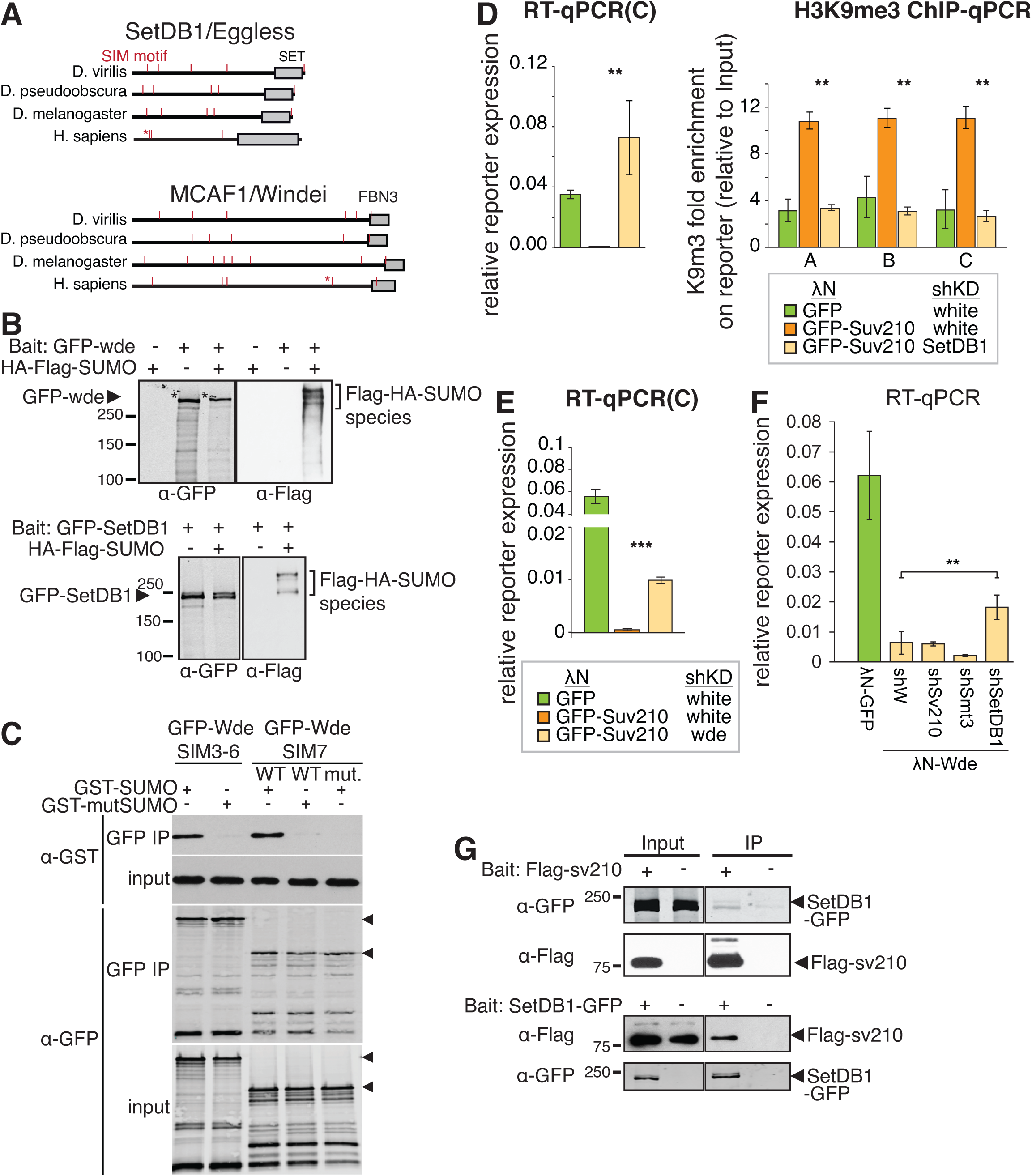
*Drosophila* SetDB1 is required for Su(var)2-10 mediated reporter silencing and physically interact with Su(var)2-10. (A)*Drosophila* SetDB1 and Wde contain predicted SIMs. Diagrams show computationally predicted SIMs (red marks) on the SetDB1 and Wde homologs in three representative *Drosophila* species and in human. The functional SIM of human SetDB1 and Wde described in Ivanov et al.(Ivanov et al., 2007) and Uchimura et al.(Uchimura et al., 2006) respectively are marked by asterisk. Grey boxes show the position of the conserved SET and FBNIII domains of the two proteins. (B)SetDB1 and Wde interact with SUMOylated proteins. Total protein lysates from S2 cells co-expressing FLAG-HA-SUMO and GFP-SetDB1 or GFP-Wde were immunopurified using anti GFP nanotrap beads. Cells not expressing FLAG-HA-SUMO were used as negative control. Western blots were probed with anti-GFP and anti-FLAG antibodies. (C)SetDB1 is required for Su(var)2-10-induced reporter repression and H3K9me3 deposition at reporter locus. Bar plots show the expression levels of reporter (RT-qPCR) and H3K9me3 occupancy at reporter locus (ChIP-qPCR at regions indicated in Fig 5A) in the ovary upon tethering of AN-GFP-Su(var)2-10 and GLKD of SetDB1 by shRNA, in comparison with AN-GFP or AN-GFP-Su(var)2-10 tethering in control background (as in Fig 5B, D). Su(var)2-10 induced reporter repression and H3K9me3 enrichment is reverted upon knockdown of SetDB1. Data is normalized to *Rp49* as internal control, and to input for ChI P experiments. Error bars show the standard deviation of three biological replicates for RT-qPCR, or two biological replicates of ChIP-qPCR. P-values for the difference between shSetDB1 and control (shW) upon AN-GFP-Su(var)2-10 tethering were determined by Student’s t-test. *p<0.05, **p<0.01. (D)The SetDB1 cofactor Wde is required for Su(var)2-10 induced reporter mRNA repression. Bar plot shows reporter expression levels (RT-qPCR) in ovary upon tethering of AN-GFP-Su(var)2-10 and GLKD of Wde by shRNA, in comparison with AN-GFP or AN-GFP-Su(var)2-10 tethering in control background. Error bars show standard deviation of three biological replicates. P-values for the difference between shWde and control (shW) upon AN-GFP-Su(var)2-10 tethering were determined by Student’s t-test; ***p<0.001. (E)Su(var)2-10 and SUMO are not required for Wde-induced repression. Boxplot shows reporter expression levels (region B) in ovaries expressing AN-GFP control or AN-GFP-Wde in control ovaries (shW) and ovaries depleted from Su(var)2-10, SUMO, and SetDB1, as estimated by RT-qPCR. Error bars show the standard deviation from 3 biological replicates. Statistical significance is estimated by one-way ANOVA followed by Tukey’s test; **p<0.01. (G)Su(var)2-10 interacts with SetDB1. Protein lysates from ovaries expressing FLAG-Su(var)2-10 and GFP-SetDB1 were immunoprecipitated with either FLAG or GFP nanotrap beads. In each experiment, lysate from ovaries not expressing the bait protein was used as negative control. Western blots were probed with anti-FLAG antibody to detect Su(var)2-10 and anti-GFP antibody to detect SetDB1.

To define the place of Su(var)2-10 in the Piwi silencing pathway, we tested if Su(var)2-10 induced repression is dependent on SetDB1 and Wde. We tethered Su(var)2-10 to the reporter and depleted SetDB1 or Wde in germ cells through shRNA-mediated knockdown. SetDB1 knockdown abolished the ability of Su(var)2-10 to silence reporter transcription and induce H3K9me3 deposition (Fig. 7D). Similarly, knockdown of the SetDB1 co-factor Wde resulted in partial release of reporter silencing (Fig. 7E). Thus, the transcriptional repression caused by recruitment of Su(var)2-10 to a genomic locus depends on H3K9me3 deposition by the histone methyltransferase SetDB1 and its co-factor Wde.

As SetDB1 and Wde have SUMO-interaction motifs, SUMOylation of chromatin-associated proteins by Su(var)2-10, including SUMOylation of Su(var)2-10 itself, might promote recruitment of the SetDB1/Wde complex to target loci. Alternatively, or in addition, interaction with SUMOylated proteins might enhance the histone methyltransferase activity of SetDB1/Wde. To test the latter possibility, we decided to tether Wde to the reporter locus in a SUMO-independent manner, and probe the involvement of Su(var)2-10 in its silencing activity. Tethering of Wde induced strong reporter repression in germ cells of the fly ovary (Fig. 7F). Importantly, this repression was dependent on SetDB1, but independent of Su(var)2-10 and SUMO, as depletion of either of these proteins did not release reporter silencing. This result suggests that SUMO and Su(var)2-10 do not impact the enzymatic activity of SetDB1/Wde and are instead involved in its recruitment to genomic targets. Co-immunoprecipitation from ovarian lysate showed that Su(var)2-10 and SetDB1 interact *in vivo* (Fig. 7G). Taken together, our results suggest that SUMOylation of protein targets by Su(var)2-10 - including SUMOylation of Su(var)2-10 itself - provides a binding platform for the recruitment of the SetDB1/Wde complex to genomic targets in order to induce H3K9 trimethylation and transcriptional repression.

## Discussion

The classic RNAi mechanism uses small RNAs to guide the endonuclease activity of Argonaute proteins for sequence-specific degradation of mRNA targets in cytoplasm. A distinct clade of Argonautes, Piwi proteins, also use this mechanism to destroy mRNA targets, however, both flies and mammals also utilize nuclear-localized Piwi proteins (Piwi in Drosophila and Miwi2 in mouse) to induce transcriptional repression. Indirect evidence suggests that the piRNA/Piwi complex recognizes its nuclear targets using complementary interactions between piRNA and nascent chromatin-bound transcripts (Pezic et al., 2014; Sienski et al., 2012). In both insect and mammals, piRNA-guided transcriptional silencing is associated with the deposition of repressive histone marks on genomic targets(LeThomas et al., 2013; Pezic et al., 2014; Rozhkov et al., 2013; Sienski et al., 2012). Studies indicated that in *Drosophila* the conserved histone methyltransferase SetDB1 (Egg) is responsible for deposition of silencing mark at Piwi targets(Sienski et al., 2015; Yu et al., 2015). However, the molecular mechanism leading to the recruitment of SetDB1 by the Piwi/piRNA complex remained unknown. Here, we showed that SUMO and the SUMO E3 ligase Su(var)2-10 are required for piRNA-guided transcriptional silencing in *Drosophila.* Several lines of evidence indicate that SUMO and Su(var)2-10 act together downstream of the piRNA-guided complex. First, depletion of SUMO and Su(var)2-10 decrease the H3K9me3 silencing mark and release TE repression induced by the piRNA pathway (Fig. 2,3,6). Second, localization of Su(var)2-10 to distinct nuclear foci depends on Piwi, Arx and Panx (Fig. 4C) and these 3 factors also co-immunoprecipitated with Su(var)2-10 (Fig. 4B), indicating genetic and physical interactions between piRNA/Piwi/Arx/Panx and Su(var)2-10. Third, SUMO and Su(var)2-10 are required for transcription silencing of reporter in an established model of repression caused by tethering of Panx to chromatin(Sienski et al., 2015; Yu et al., 2015) (Fig. 4A). Finally, recruitment of Su(var)2-10 to artificial targets on chromatin leads to transcriptional repression accompanied with SUMO accumulation and H3K9me3 deposition at the target locus (Fig. 5H-J). Repression induced by SUMO and Su(var)2-10 depends on the methyltransferase complex composed of SetDB1 and Wde, which are recruited to the target locus through their interaction with SUMO (Fig. 7C-E). Taken together, our results suggest a model for the molecular mechanism of piRNA-guided transcriptional silencing that fills the gap between the target recognition complex composed of piRNA/Piwi/Panx/Arx and the chromatin effector complex composed of SetDB1 and Wde (Fig. 8).

**Figure 8.**
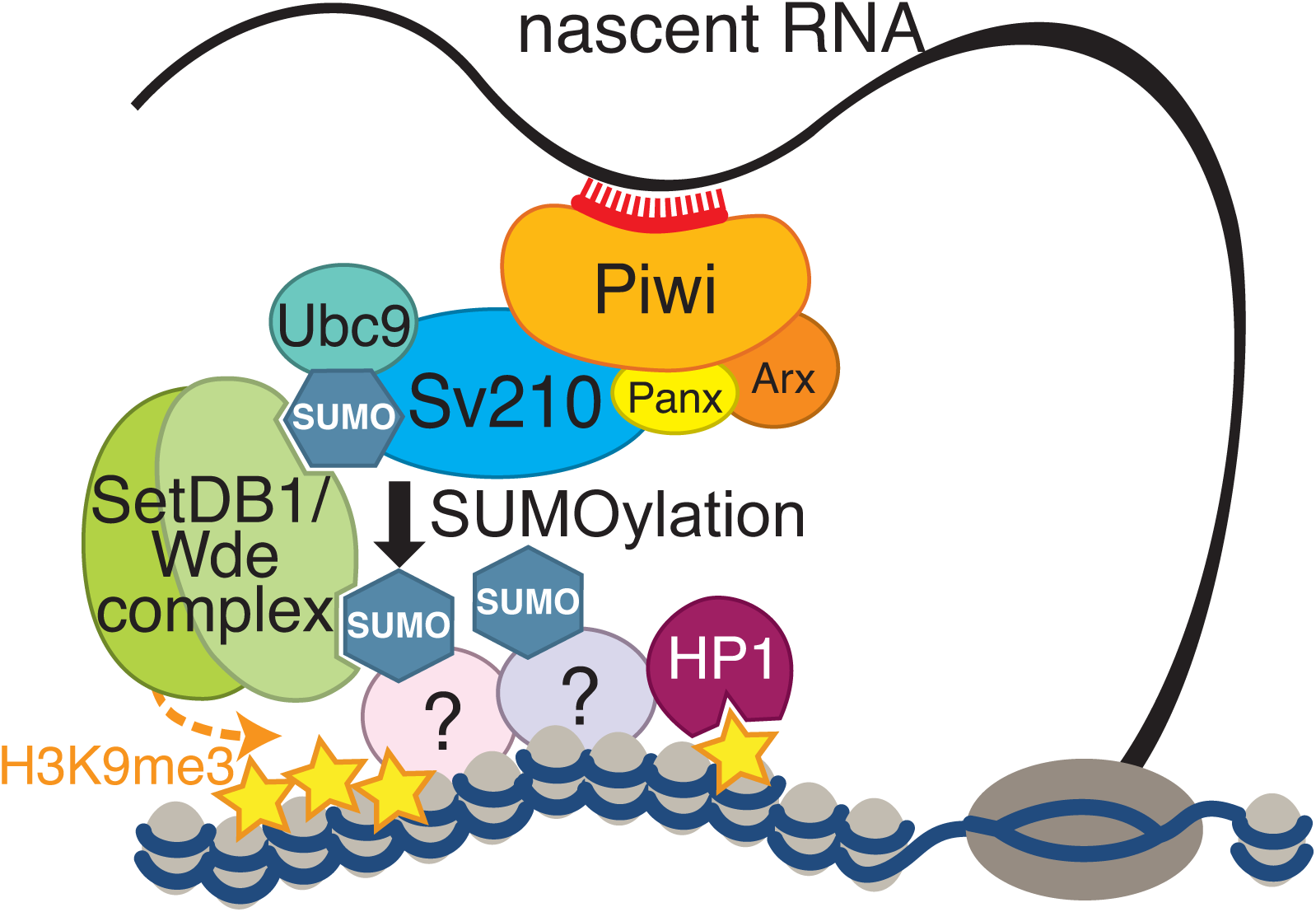
Model of piRNA-guided H3K9me3 deposition. piRNA-loaded Piwi complex recognizes nascent transcripts. Su(var)2-10 interacts with the piRITS complex and promotes SUMO modification of itself and yet-to-be identified chromatin factor(s). The SUMO moiety of Su(var)2-10 and chromatin factors recruits the SetDB1/Wde complex through their SIM interaction motifs, which installs the silencing H3K9me3 mark at target loci.

Our results identified a new role for the SUMO pathway in piRNA-guided transcriptional silencing. Recently, the SUMO pathway emerged as an important factor in heterochromatin maintenance and genome stability in different organisms. SUMOylation through PIAS enzymes has a conserved role in repair of double strand DNA breaks in yeast and *Drosophila* (Psakhye and Jentsch, 2012; Ryu et al., 2015). The SUMO modification was demonstrated to act as a binding platform for heterochromatin effectors such as HP1 in various systems from yeast to humans(Ivanov et al., 2007; Maison et al., 2011, 2016; Shin et al., 2005; Stielow et al., 2008b; Thompson et al., 2015; Uchimura et al., 2006), suggesting that SUMO has a conserved role in heterochromatin assembly. Importantly, in mammalian systems SUMO is required for recruitment of the histone methyltransferase complex composed of SetDB1 and MCAF1 to its chromatin targets and for its activity(Ivanov et al., 2007; Stielow et al., 2008b; Thompson et al., 2015; Uchimura et al., 2006). SetDB1 and MCAF1 (called Wde in Drosophila) are conserved between insects and mammals. We found that Drosophila SetDB1 and Wde contain functional SUMO-interaction motifs (Fig 7A, C). Furthermore, SetDB1-dependent repression induced by tethering of Su(var)2-10 to the reporter requires SUMO (Fig 6A) and correlates with increased SUMO ChIP signal (Fig 5H).

Remarkably, the SUMO-dependent recruitment of SetDB1 is involved in silencing of transposable elements in somatic cells in mammals, although the molecular mechanism of this process is different and does not require piRNA. Instead, members of the large vertebrate-specific family of Krüppel-associated box domain-zinc finger proteins (KRAB-ZFPs) repress endogenous retroviruses through binding specific DNA motifs in their sequences (reviewed in (Wolf et al., 2015)). Although distinct members of the KRAB-ZFP family recognize different sequence motifs in target transposons, repression of all targets by various KRAB-ZFPs requires the universal co-repressor KAP1/TIF1b (KRAB-associated protein 1). KAP1 has a SUMO E3 ligase activity and its auto-SUMOylation leads to recruitment of SetDB1(Ivanov et al., 2007). Our results suggest that *Drosophila* Su(var)2-10 can be SUMOylated (Fig S4C), and SetDB1 and Wde have functional SUMO-interacting motifs (SIMs) (Fig 7), suggesting that auto-SUMOylation of Su(var)2-10 might induce the recruitment of the SetDB1/Wde complex (Fig 8). Transposon repression by DNA-binding proteins (KRAB-ZFPs) and the piRNA pathway are two conceptually different mechanisms of target recognition, but both mechanisms lead to SUMO-dependent recruitment of conserved chromatin-modifying activity to the target. Repression by KRAB-ZFPs requires large family of DNA-binding proteins with each member designed to recognize specific target. In contrast, the piRNA pathway employs an elegant approach of utilizing a single Piwi protein that can be programmed to target any sequence by loading different small RNA guides. The next step - recruitment of a SUMO E3 ligase by the target-recognition complex - is similar in the two pathways, however, it should be noted that even though both KAP1 and Su(var)2-10 have SUMO ligase activity, these two proteins are not homologous, suggesting convergent evolution of the two pathways. The final step of transposon repression - SUMO-dependent recruitment of the histone methyltransferase complex and installation of the repressive chromatin marks - seems to be identical in the two pathways.

The nuclear Piwi proteins in *Drosophila* and mouse, PIWI and MIWI2, are not one-to-one orthologs. Unlike *Drosophila*, other insects including the silkworm *Bombyx mori*, the flour beetle *Tribolium castaneum* and the honeybee *Apis mellifera* encode only two Piwi proteins and, at least in *B. mori*, these proteins do not localize to the nucleus (Nishida et al., 2015). These observations suggest that the nuclear Piwi pathway in *Drosophila* has evolved independently in this lineage. In light of this evolutionary interpretation, the interaction of the Piwi complex and the E3 SUMO ligase Su(var)2-10 indicates that the nuclear piRNA pathway co-opted an ancient mechanism of SUMO-dependent recruitment of the histone modification complex for transcriptional silencing of transposons in *Drosophila.* The molecular mechanism of piRNA-induced transcriptional repression in other clades such as mammals might have evolved independently of the corresponding pathway in flies. It will be interesting to investigate if mammals also use SUMO-dependent recruitment of chromatin repressor complexes for transcriptional repression of piRNA targets.

Our results and studies in mammals(Ivanov et al., 2007) suggest that in both clades self-SUMOylation of SUMO E3 ligases might be involved in recruitment of the SetDB1 histone methyltransferase complex to chromatin. However, these results do not exclude the possibility that the recruitment of SetDB1 is facilitated by SUMOylation of additional chromatin proteins by Su(var)2-10. Studies in yeast led to the proposal of the ‘SUMO spray’ hypothesis that postulates that SUMOylation of multiple different proteins localized in physical proximity promotes the assembly of multi-unit effector complexes(Psakhye and Jentsch, 2012). Local concentration of multiple SUMO moieties leads to efficient recruitment of SUMO-interacting proteins. According to this hypothesis multiple SUMO-SIM interactions within a protein complex act synergistically, thus SUMOylation of any single protein is neither necessary nor sufficient to trigger downstream processes(Jentsch and Psakhye, 2013; Psakhye and Jentsch, 2012). Assembly of such ‘SUMO spray’ on chromatin might be governed by the same principles of multiple weak interactions as was recently recognized for the formation of various phase-separated liquid-droplet compartments in the cell (Shin and Brangwynne, 2017). The presence of Su(var)2-10 on a chromatin locus might lead to SUMOylation of multiple chromatin-associated proteins that are collectively required for the recruitment of effector chromatin modifiers. The SUMOylation consensus (ψKxE/D) is very simple and therefore quite common in the fly proteome. Consistent with this, several hundred SUMOylated proteins were identified in proteomic studies in Drosophila(Handu et al., 2015; Nie et al., 2009). SetDB1 was shown to be a target of SUMOylation and it was proposed that SetDB1 SUMOylation may facilitate the formation of multi-subunit chromatin-modifying complexes(Koch et al., 2009). Thus, it is possible that collective SUMOylation of multiple chromatin-associated proteins contributes to recruitment and stabilization of the SetDB1 complex on chromatin (Fig. 8).

The cascade of events leading to repression initiated by target recognition by piRNA/Piwi, followed by interaction with Su(var)2-10 and subsequent SUMO-dependent recruitment of SetDB1/Wde suggests that the three complexes tightly cooperate. But do these three complexes (Piwi, Su(var)2-10 and SetDB1) always work together, or does each complex have additional functions independent of the other two? Genome-wide analysis suggests that the vast majority of targets of piRNA-guided transcriptional silencing are repressed through SUMO/Su(var)2-10 and, likely, SetDB1/Wde, suggesting that Piwi always requires these other complexes for its function in transcriptional silencing. However, our data and published studies suggest that SetDB1 can be recruited to some genomic targets in a manner that is independent of both piRNAs and Su(var)2-10. For example, in the accompanying paper we describe host genes targeted by Su(var)2-10 that are repressed through SUMO- and SetDB1-dependent H3K9 trimethylation but independently of piRNAs (Ninova et. al, accompanying manuscript). Su(var)2-10 and SetDB1 are also expressed outside of the gonads and were implicated in chromatin silencing in somatic tissues that lack an active piRNA pathway (Brower-Toland et al., 2009; Hari et al., 2001; Seum et al., 2007; Stampfel et al., 2015; Stielow et al., 2008a; Tzeng et al., 2007). Our studies reveal that cooperation between Su(var)2-10 and SetDB1 is important for tissue-specific restriction of gene expression and maintenance of homeostatic balance between eu- and heterochromatin (see Ninova et al., accompanying manuscript).

## MATERIALS AND METHODS

### Fly stocks and eggshell phenotyping

The stocks for shRNAs of *Su(var)2-10* (shSv210-1 and shSv210-2, BDSC #32915 and BDSC #32956, respectively), *piwi* (shPiwi, BDSC #33724), *wde* (shWde, BDSC BL33339) and *white* (shWhite, BDSC #33623) were obtained from the Bloomington Drosophila Stock Center. UASp-mKate2-4xBoxB-K10polyA, UASp-λN-GFP-eGFP control, GFP-Piwi, GFP-Arx and shPiwi were described previously(Chen et al., 2016; LeThomas et al., 2013). shSetDB1, shPanx, λN -Panx and Tubulin-BoxB reporter stocks were gifts from Julius Brennecke, the luciferase 10BoxB reporter is a gift from Gregory Hannon. To obtain the shAsterix line, the short hairpin sequence was ligated into the pValium20 vector (Ni et al. 2011) using T4 DNA ligase from NEB (M0202), according to the manual, and then integrated into the attP2 landing site (BDSC #25710). Hairpin sequences are listed in Supplementary Table S1. shSmt3 (shSUMO) was reconstructed based on the TRiP line HMS01540 and integrated into the attP2 landing site (BDSC #8622). For all other *Drosophila* lines generated in this study, respective full length cDNA sequences or mutants were cloned in pENTR™ /D-TOPO® (Invitrogen) entry vectors and transferred to Gateway® destination vectors containing attB site, a miniwhite marker followed by UASp promoter sequence, and λN-GFP upstream the gateway cassette, or GFP downstream the gateway cassette. Transgenic flies carrying these constructs were generated by phiC31 transformation at BestGene Inc. UASp-SetDB1-GFP was integrated into attP-3B landing site (BDSC #9750). UASp-λN-GFP-Su(var)2-10-PA was integrated into attP9A landing site (BDSC #9736). UASp-λN-GFP-Panx, UASp-λN-GFP-Arx, UASp-FLAG-Su(var)2-10, UASp-λN-GFP-Su(var)2-10-C341S, UASp-λN-GFP-Su(var)2-10-ΔSAP, UASp-λN-GFP-Su(var)2-10-ΔPINIT, UASp-λN-GFP-Su(var)2-10-ΔSP-RING were integrated into the attP40 landing site (y^1^ w^67c23^; P{CaryP}attP40).

The expression of all constructs was driven by maternal alpha-tubulin67C-Gal4 (MT-Gal4) (BDSC #7063 or #7062), except for the experiment of Su(var)2-10 depletion in the ovarian soma where Tj-gal4 (DGRC #104055) driver was used. All flies were put on yeast for 2 to 3 days before ovary dissections and were at the age of 3 to 14 days after hatching. For eggshell phenotyping, freshly laid eggs were mounted in 1XPBS and manually counted under a dissecting microscope.

### Imaging

Ovaries from *Drosophila* lines expressing UASp-λN-GFP-Su(var)2-10 and UASp-driven shRNAs against *white, Arx, Panx* and *Piwi* under the control of the maternal-tubulin-Gal4 driver were fixed in PBS supplemented with 4% formaldehyde for 20 minutes at room temperature with end-to-end rotation. Samples were washed three times 10 minutes with PBS, and mounted in ProLong Gold Antifade Mountant with DAPI. Imaging was performed using a Zeiss LSM 880 confocal microscope and data was processed using Fiji(Schindelin et al., 2012).

### RNA extraction and RT-qPCR

RNA was isolated with Ribozol (Amresco, N580) and treated with DNaseI (Invitrogen, 18068-015). Reverse transcription was carried out using Superscript III (Invitrogen) with random hexamers. qPCR was performed on a Mastercycler®ep realplex PCR thermal cycler machine (Eppendorf). Primers used in qPCR are listed in Supplementary Table 1.

### Immunoprecipitation and western blot from ovaries

For immunoprecipitation (IP), 50-70 pairs of freshly dissected ovaries were lysed with a douncer in 500μl lysis buffer (0.2% NP40, 20 mM Tris pH7.4, 150 mM NaCl, 10% glycerol) supplied with protease inhibitor (Roche, 11836170001) and 20 mM deSUMOylation inhibitor N-Ethylmaleimide (Sigma, E3876). For Piwi/Arx/Panx coIP with Su(var)2-10 the lysis buffer contained 0.4% NP40. For GFP IP, lysates were incubated with GFP-Trap® or control (ChromoTek) magnetic agarose beads for 1-2 hr at 4°C with end-to-end rotation. For FLAG IP, lysates were incubated with anti-FLAG M2 ® magnetic beads (Sigma M8823). After incubation, the beads were washed 5 times with 500μl wash buffer (0.1% NP40, 20 mM Tris pH7.4, 150mM NaCl) containing protease inhibitor and 20 mM N-Ethylmaleimide. For Piwi/Arx/Panx coIP with Su(var)2-10 the wash buffer contained 250mM salt and 0.2% NP40. The washed beads were boiled in 75 μl SDS-PAGE sample buffer, and then the supernatant was used for western blot analysis. Western blots were carried out using the following antibodies: anti-FLAG [Sigma, A8592], anti-GFP [ab290], or rabbit polyclonal anti-GFP(Chen et al., 2016), and HRP-conjugated or IRDye® anti-rabbit and anti-mouse secondary antibodies (Li-cor #925-68070 and −68071, 925-32210 and −32211, Cell Signalling 7074, 7076).

### Immunoprecipitation from S2 cells

S2 cells were cultured in Schneider’s *Drosophila* Medium containing 10% heat-inactivated FBS and 1X Penicillin-Streptomycin. Expression vectors encoding GFP-fusion and FLAG-fusion proteins under the control of the Actin promoter were generated from entry cDNA clones transferred to pAGW or pAFW destination vectors from the *Drosophila Gateway*™ *Vector* collection using the Gateway® system. GFP-wde expression vector was a generous gift from Andreas Wodarz (Koch et al., 2009). S2 cells were transfected with T ransIT-LT1 (Mirus). 24-48h post transfection cells were harvested and lysed in lysis buffer (20 mM Tris-HCl at pH 7.4, 150 mM NaCl, 0.2% NP-40, 0.2% Triton-X, 5% glycerol), supplemented with protease inhibitor cocktail (Roche, 11836170001) and 20mM N-Ethylmaleimide. Co-immunoprecipitation experiments and western blots were performed as described for ovarian tissue.

### GST-SUMO interaction assays

pGEX-2TK vectors (GE Healthcare) expressing GST-SMT3 (SUMO) (plasmid was a generous gift from G Suske (Stielow et al., 2008a)) and SIM interaction deficient GST-SMT3 generated in our lab were transformed in *E. coli* strain BL21 and purified by glutathione affinity chromatography using a standard protocol. In brief, Glutathione Sepharose 4B (GE Healthcare) slurry was equilibrated using five bead volumes of 50 mM Tris-HCl pH 7.4, 150 mM NaCl. Bacteria were lysed in 50 mM Tris-HCl pH 7.4, 150 mM NaCl, 1 mM buffer using a French press, and slurry was incubated with bacterial lysate at 4°C with end-over-end rotation. After three washes with the same buffer, the fusion proteins were eluted in a buffer containing 50 mM Tris HCl (pH 8.0), 150 mM NaCl and 20 mM reduced glutathione. Eluates were dialysed with 10 kDa cut-off dialysis tubing against Tris-HCl buffer (50 mM Tris-HCl, pH 7,5, 500 mM NaCl and 1 mM DTT) overnight at 4°C.

S2 cells transfected with plasmid encoding either GFP-Su(var)2-10-PA, GFP-tagged truncated Wde including predicted SIM-s 3 to 6 (aa385-655), GFP-tagged truncated Wde including SIM 7 (aa1058-1310), or GFP-tagged truncated Wde with mutated SIM 7 (1202DL>AA) (see Figure S5 for Wde map and SIM motif annotations) under the control of pActin promoter. Cells were harvested 24-48h post transfection.

For Su(var)2-10-SUMO interaction, cells were lysed in RIPA buffer supplemented with protease inhibitor (Roche 11836170001) and 20 mM NEM. Cleared lysates were diluted 1:10 with binding buffer (20 mM Tris-HCl, pH 7.4, 150 mM NaCl, 0.1% NP40) and pre-cleared by rotation with Glutathione Sepharose 4B slurry (GE Healthcare) for 1 hour at 4°C with end-to-end rotation. Lysates were divided in equal parts and incubated with 2 μg purified GST-SUMO(wild type), GST-SUMO(mutant) and 10 μl Glutathione Sepharose 4B slurry for 2hours at 4°C with end-to-end rotation. Beads were washed 4 times for 10 minutes with binding buffer and boiled in SDS-PAGE protein loading buffer. Bound proteins were analysed by western blot using anti-GFP antibody.

For Wde-SUMO interactions, cells were lysed in lysis buffer (20 mM Tris-HCl at pH 7.4, 150 mM NaCl, 0.2% NP-40, 0.1% Triton-X, 5% glycerol), supplemented with protease inhibitor (Roche 11836170001) and 20 mM NEM. Cleared lysates were incubated with 2-3 μg GST-SUMO for 2 hr, at 4°C with end-to-end rotation. Next, lysates were incubated with GFP Nanotrap beads (Chromotek), pre-blocked with 0.5% BSA and untransfected S2 cell lysates, for 1 hr at 4°C with end-to-end rotation. Beads were washed 5 times for 10 minutes at 4°C with wash buffer (0.1% NP-40, 20 mM Tris-HCl pH 7.4, 150 mM NaCl) and finally boiled in SDS-PAGE protein loading buffer. Bound proteins were analysed by western blot using anti-GFP antibody (ab290, Abcam) and anti-GST antibody (Cell Signalling #2622).

### In vitro SUMOylation assay

His-Su(var)2-10 was cloned into the pGEX-2TK vector and expressed in *E. coli* strain BL21. Protein was purified using His-Pur Ni-NTA Resin (Thermo Scientific), equilibrated using 2 bed volumes of PBS pH=7.4. The fusion protein was eluted in a buffer containing PBS pH=7.4 and 250 mM imidazole. The elution fractions were dialysed using 10 kDa cut-off dialysis tubing in PBS supplemented with 1 mM DTT overnight at 4°C. After dialysis protein was concentrated using 10kDa MWCO Amicon filter (Millipore). *In vitro* SUMOylation assay was performed using SUMO2 Conjugation Kit (BostonBiochem, K-715) according to the manufacturer’s instruction, using 5μl 34μM stock of recombinant Su(var)2-10. Reaction without ATP was set up as a negative control. Results were detected using Western Blotting using anti-His primary antibody (ThermoFisher, His.H8).

### RNA-seq

For RNA-seq, ovarian total RNA (10-15 μg) from shW, shSv210-1, shSv210-2, and shSmt3 GLKD lines was depleted of ribosomal RNA with the Ribo-Zero™ rRNA Removal Kit (Epicentre/Illumina). Initial RNA-seq libraries were made using the TruSeq RNA prep kit by Illumina (shW, shSv210-1, shSv210-2). A second set of libraries from shW, shSv210-2 and shSmt3 in two biological replicates were made using the NEBNext® Ultra™ Directional RNA Library Prep Kit. Libraries were sequenced on the Illumina HiSeq 2000/2500 platform. RNA-seq data from shPiwi and corresponding shW control lines were previously published (LeThomas et al., 2013).

### ChIP-seq and ChIP-qPCR

ChIPs were carried out as described previously(LeThomas et al., 2014) with the following antibodies: anti-H3K9me3 [ab8898], anti-RNA Pol II [ab5408], anti-H3K4me2/3 [Ab6000], anti-H3K36me3 [ab9050], HP1a [C1A9, DSHB] and *anti-Drosophila* SUMO (smt3), a kind gift from G Cavalli(Gonzalez et al., 2014). ChIP-seq library construction was carried out using the NEBNext ChIP-Seq Library Prep Master Mix Set (E6240) with minor modifications. After adaptor ligation and PCR amplification, size selections were done on a 2% agarose gel to select the 200bp-400bp size window. Libraries were sequenced on the Illumina HiSeq 2000/2500 platform (SR 49bp or 50bp). ChIP-qPCR was performed on a Mastercycler®ep realplex PCR thermal cycler machine (Eppendorf). Primers used in ChIP-qPCR are listed in Table 1. All ChIPs were normalized to respective inputs and to control region *rp49.* H3K9me3 ChIP-seq data from shPiwi and corresponding shW control lines were previously published(LeThomas et al., 2013).

### Bioinformatic analyses

The *D. melanogaster* genome assembly BDGP RP/dm3 (April 2006) was used in all analysis. All alignments were performed using an in-house pipeline employing Bowtie 0.12.17(Langmead et al., 2009).

RNA-seq datasets were pre-processed to remove adaptor contamination and rRNA sequences, as described previously(Chen et al., 2016). The remaining reads were aligned to the *D. melanogaster* genome allowing 2 mismatches, and retaining only uniquely mapping reads. For analysis of transposons, data were mapped allowing up to 10,000 mapping positions and 0 mismatches, and read counts were corrected based on the number of mapped positions as described previously(Manakov et al., 2015). Protein-coding gene annotations and TE annotations were obtained from the RefSeq and RepeatMasker tables, respectively, retrieved from the UCSC genome browser(Karolchik et al., 2004). TE expression values for TE families were calculated as the sum of mappability-corrected reads aligning to individual repeats. Similar results were obtained by an alternative approach were RNA-seq reads were directly aligned to RepeatMasker TE consensuses allowing 3mismatches (data not shown). For scatter plots, read counts per element were normalized as reads per million mapped reads (RPKM). Differential expression analyses were performed using the DESeq2 R package using raw read counts as input(Love et al., 2014).

ChIP-seq data was aligned allowing 2 mismatches and retaining uniquely mapped reads for analysis of unique regions. As in RNA-seq, for TE analyses, reads were mapped allowing up to 10,000 mapping positions and 0mismatches, and read counts were corrected based on the number of mapped positions. Read counts for 1Kb genomic windows or individual TEs were normalized as RPKM. ChIP signal was defined as the ratio of normalized ChIP to Input counts (ChIP/Input). For global analysis, 1Kb genomic windows that had less than 1RPKM in input libraries were excluded. 1Kb genomic windows with ChIP/Input ratio > 2 in all control ChIP-seq datasets (shW) were annotated as “het”. We note that using higher ChIP/Input ratio cutoffs to define heterochromatic windows (stricter H3K9me3 enrichment cutoff) produces similar results (data not shown). Heatmaps were generated using the ‘pheatmap’ R package. Normalized genome coverage tracks were generated using BedTools(Quinlan and Hall, 2010) and BigWig tools(Kent et al., 2010), using the total mapped reads as a scaling factor. Circular plot was generated using Circos 0.67.7(Krzywinski et al., 2009). Non-reference insertions were annotated using the TIDAL pipeline(Rahman et al., 2015) with default parameters, using merged reads that do not map to the reference genome with 2 mismatches from all experiments involving DNA sequencing (DNA from Input and ChIPs).

Orthologs of SetDB1 and Wde in *Drosophila* species were extracted from OrthoDB1(Waterhouse et al., 2011) and UniProt databases. SIM predictions were performed using the GPS-SUMO online tool(Zhao et al., 2014). Protein sequences were aligned by MUSCLE(Edgar, 2004) and visualized in Jalview (Clamp et al., 2004).

### Data availability

RNA-seq and H3K9me3 ChIP-seq data from shPiwi and corresponding shW control lines were previously published(LeThomas et al., 2013). All RNA-seq and ChIP-seq data generated in this study is deposited to the Gene Expression Omnibus database: GSE115277.

## Acknowledgements

We thank members of the Fejes Toth and Aravin labs for discussion. We thank Gary Karpen for suggestions and discussion of some of the experiments. We appreciate the help of Kathy Situ, Zsófia Török, Sivani Vempati, Solomiia Khomandiak, and Angel Galvez Merchan with the experiments. We are grateful to Julius Brennecke, Gregory Hannon and the Bloomington Stock Center for providing fly stocks, Julius Brennecke and Giacomo Cavalli for providing antibodies, Andreas Wodarz for the GFP-wde expression vector, and Guntram Suske for the GST-Smt3 (wild type) plasmid. We thank Igor Antoshechkin (Caltech) for help with sequencing and Sergei Manakov for bioinformatic support. This work was supported by grants from the National Institutes of Health (R01 GM097363), the Ministry of Education and Science of Russian Federation (14.W03.31.0007) and by the Packard Fellowship Awards to A.A.A., and the National Institutes of Health (R01GM110217) and Ellison Medical Foundation Awards to K.F.T. AKR was an NSF GRFP fellow.

## Author contributions

MN, YAC, AAA and KFT designed the experiments. MN, YAC, BG, AR, KFT and YL executed the experiments. MN performed the computational analysis and interpretation of the data. The manuscript was written by MN, YAC, AAA and KFT.

## Declaration of Interests

The authors declare no competing interests.

## Supplementary Figure Legends

**Supplementary Figure S1. Expression of transposons and protein-coding genes upon Su(var)2-10 knockdown**

(A)TE upregulation upon germline knockdown of Su(var)2-10 using the Su(var)2-10-1 hairpin. Scatter plot shows log2-tansformed RPKM values for TEs (RepeatMasker) in RNA-seq data from ovaries expressing shSv210-1 (Y-axis) versus shW control hairpin (X-axis) under the control of the MT-gal4 driver. Dashed lines indicate 4-fold change.

(B)TE upregulation is reproducible upon Su(var)2-10 knockdown in the ovary. Heatmaps reflect the expression levels (RPKM normalized) of TE families in two biological replicates of control (shW) and Su(var)2-

10knockdown, using the stronger Su(var)2-10 hairpin (shSv210-2, see main text). Coloured bars correspond to RepeatMasker annotated transposon families. Data was sorted by Euclidean distance.

(C)(Left) Scatter plot showing the average expression (X-axis) versus fold change in Su(var)2-10-2 KD (shSv210-2) compared to control (shW) ovaries (Y-axis) for protein-coding genes and transposons in RNA-seq data. Significantly changed protein-coding genes and TEs (5% FDR cutoff) are marked. Analysis was performed with the DESeq2 R-package(Love et al., 2014) with 2 biological replicates for each condition.

(Right) Boxplot showing the distribution of RNA-seq fold changes between Su(var)2-10 KD and control (shW) ovaries for protein-coding (PC) genes, and RepeatMasker transposons (TEs RMSK).

**Supplementary Figure S2. Distribution of repetitive sequences on H3K9me3-enriched regions.**

Boxplots show the proportion of RepeatMasker annotations in the sequence of 1kb genomic intervals which have >2-fold enrichment of H3K9me3 signal (“het”) and H3K9me3-poor intervals (“eu”).

**Supplementary Figure S3. Genetic and biochemical interaction of Su(var)2-10 with Panx, Arx and Piwi**

(A) Su(var)2-10 interacts with Arx in S2 cells. Lysates from S2 ectopically expressing GFP-Su(var)2-10 and FLAG-tagged Arx or FLAG-mKate (negative control) were analysed by immunoprecipitation with anti-FLAG antibody. Western blot was probed with anti-FLAG antibody to detect Arx, and anti-GFP antibody to detect Su(var)2-10.

(B, C) Su(var)2-10 interacts with Piwi in S2 cells. Lysate from S2 ectopically expressing FLAG-tagged Piwi(B) or Panx(C) and GFP-Su(var)2-10 were analysed by immunoprecipitation with anti-GFP nanobody. Cells that do not express the bait protein, GFP-Su(var)2-10, were used as negative control. Western blot was probed with anti-FLAG antibody to detect Piwi/Panx, and with anti-GFP antibody to detect Su(var)2-10.

**Supplementary Figure S4.**

(A)Su(var)2-10 tethering to a Tubulin-BoxB reporter incudes transcriptional repression and H3K9me3 deposition. Bar plots show the expression level (RT-qPCR) and enrichment of H3K9me3 (ChIP-qPCR) at the Tubulin-BoxB reporter upon tethering of λN-Su(var)2-10 or λN-GFP. Data is normalized to Rp49 and, in ChIP-qPCR to input. Error bars show the standard deviation of three biological replicates for RT-qPCR, or two biological replicates of ChIP-qPCR.

(B)Su(var)2-10 interacts with SUMO *in vitro.* Bacterially purified His-tagged Su(var)2-10 was incubated with GST-SUMO(wild type) or interaction-deficient mutant GST-SUMO (QFI>AAA). GST-SUMO was affinity purified using glutathione sepharose beads and His-Su(var)2-10 was detected by Western blotting using anti-His antibody.

(C)Su(var)2-10 SUMOylation *in vitro.* Bacterially purified His-Su(var)2-10 is incubated with SUMO, SUMO E1 and E2 ligases in the presence or absence of ATP. Western blot was probed with anti-His antibody.

(D)The role of Su(var)2-10 domains in reporter silencing. λN-GFP, λN-GFP-Su(var)2-10 (wild type) or λN-GFP-Su(var)2-10 with deletions of the entire SAP, PINIT or SP-RING domain (see main text) were recruited to the mKate-4BoxB reporter. Bar plots show reporter expression in each condition as measured by RT-qPCR. Wild type and SAP-domain deletion mutant Su(var)2-10 repress the reporter, while PINIT and SP-RING domain mutants are unable to induce silencing. Error bars show standard deviation of three biological replicates Asterisks mark significant downregulation compared to the λN-GFP negative control at p<10^−6^ (one-way λNOVA followed by Tukey’s test).

**Supplementary Figure S5.**

(A)Related to Figure 7A. Multiple sequence alignments of homologs of SetDB1 and Wde in *Drosophila* species with annotated genomes. Orthologs were extracted from OrthoDB and aligned using MUSCLE. For full-length alignments, schematic of positions coloured by % identity are shown. Histograms reflect conservation. Predicted SIMs(Zhao et al., 2014) in *D. melanogaster* are marked by red bars. Regions of sequence alignments flanking the most highly conserved SIMs are magnified. Images are based on alignments edited in Jalview(Clamp et al., 2004).

(B)Su(var)2-10 interacts with SetDB1 in S2 cells. Lysate from S2 cells ectopically expressing FLAG-tagged SetDB1 and GFP-Su(var)2-10 was analysed by immunoprecipitation with anti-GFP nanobody. Cells that do not express the bait protein, GFP-Su(var)2-10, were used as negative control. Western blot was probed with anti-FLAG antibody to detect SetDB1 and anti-GFP antibody to detect Su(var)2-10.

**Supplementary Table 1.**
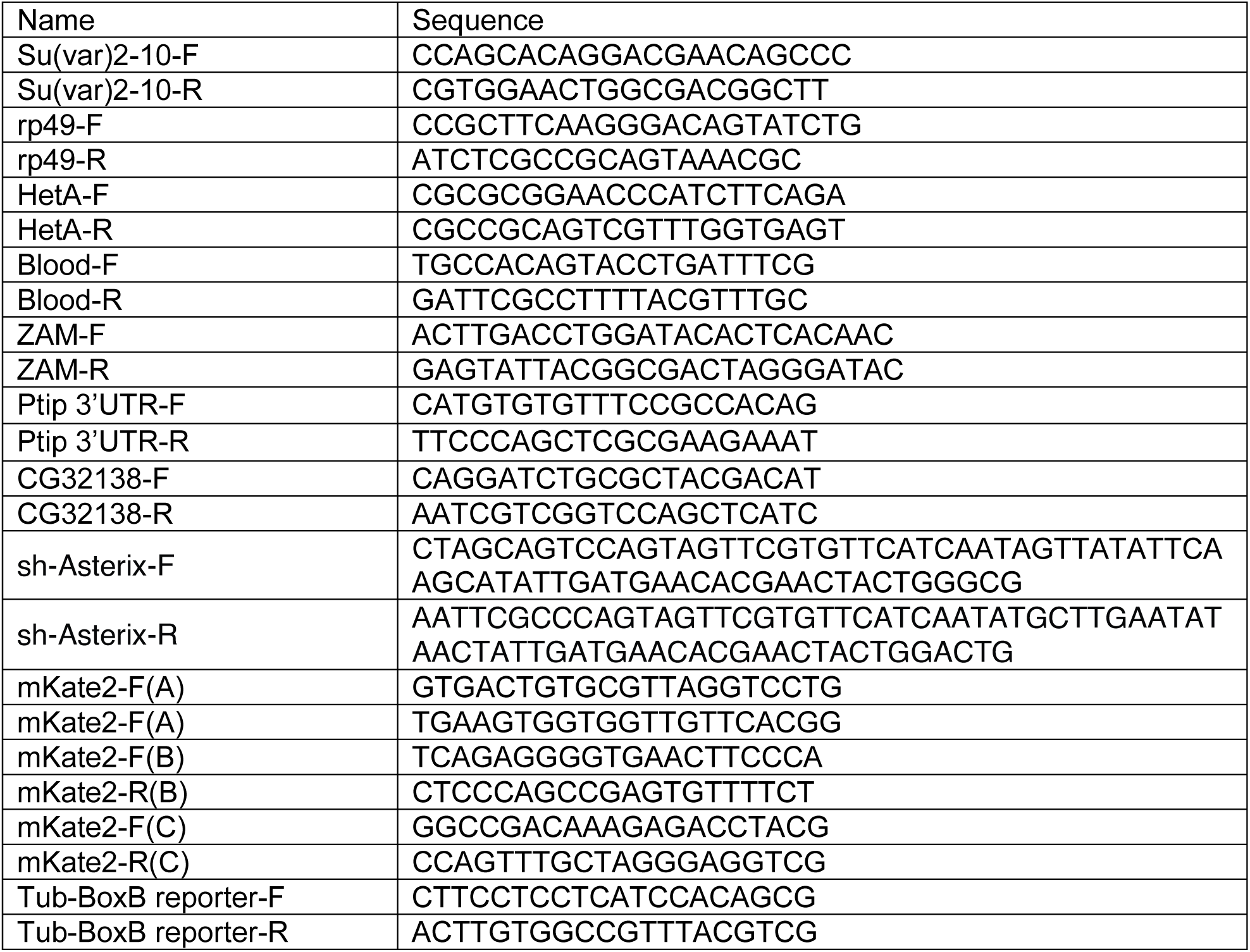
Primer list

## References

Abdu, U., Brodsky, M., and Schupbach, T. (2002). Activation of a meiotic checkpoint during Drosophila oogenesis regulates the translation of gurken through Chk2/Mnk. Curr. Biol. 12, 1645–1651.

Alekseyenko, A.A., Gorchakov, A.A., Zee, B.M., Fuchs, S.M., Kharchenko, P. V., and Kuroda, M.I. (2014). Heterochromatin-associated interactions of Drosophila HP1a with dADD1, HIPP1, and repetitive RNAs. Genes Dev. 28, 1445–1460.

Aravin, A.A., Sachidanandam, R., Bourc’his, D., Schaefer, C., Pezic, D., Toth, K.F., Bestor, T., and Hannon, G.J. (2008). A piRNA pathway primed by individual transposons is linked to De Novo DNA methylation in mice. Mol. Cell 31, 785–799.

Bannister, A.J., Zegerman, P., Partridge, J.F., Miska, E.A., Thomas, J.O., Allshire, R.C., and Kouzarides, T. (2001). Selective recognition of methylated lysine 9 on histone H3 by the HP1 chromo domain. Nature 410, 120–124.

Bernatavichute, Y. V., Zhang, X., Cokus, S., Pellegrini, M., and Jacobsen, S.E. (2008). Genome-wide association of histone H3 lysine nine methylation with CHG DNA methylation in Arabidopsis thaliana. PLoS One 3.

Betz, A., Lampen, N., Martinek, S., Young, M.W., and Darnell, J.E. (2001). A Drosophila PIAS homologue negatively regulates stat92E. Proc. Natl. Acad. Sci. 98, 9563–9568.

Brower-Toland, B., Riddle, N.C., Jiang, H., Huisinga, K.L., and Elgin, S.C.R. (2009). Multiple SET methyltransferases are required to maintain normal heterochromatin domains in the genome of Drosophila melanogaster. Genetics 181, 1303–1319.

Carmell, M.A., Girard, A., van de Kant, H.J.G.G., Bourc’his, D., Bestor, T.H., de Rooij, D.G., and Hannon, G.J. (2007). MIWI2 Is Essential for Spermatogenesis and Repression of Transposons in the Mouse Male Germline. Dev. Cell 12, 503–514.

Chen, Y.-C.A., Stuwe, E., Luo, Y., Ninova, M., Le Thomas, A., Rozhavskaya, E., Li, S., Vempati, S., Laver, J.D., Patel, D.J., et al. (2016). Cutoff Suppresses RNA Polymerase II Termination to Ensure Expression of piRNA Precursors. Mol. Cell 63, 97–109.

Chintapalli, V.R., Wang, J., and Dow, J.A.T. (2007). Using FlyAtlas to identify better Drosophila melanogaster models of human disease. Nat Genet 39, 715–720.

Chung, C.D., Liao, J., Liu, B., Rao, X., Jay, P., Berta, P., and Shuai, K. (1997). Specific inhibition of Stat3 signal transduction by PIAS3. Science 278, 1803–1805.

Clamp, M., Cuff, J., Searle, S.M., and Barton, G.J. (2004). The Jalview Java alignment editor. Bioinformatics 20, 426–427.

Donertas, D., Sienski, G., and Brennecke, J. (2013). Drosophila Gtsf1 is an essential component of the Piwi-mediated transcriptional silencing complex. Genes Dev. 27, 1693–1705.

Edgar, R.C. (2004). MUSCLE: multiple sequence alignment with high accuracy and high throughput. Nucleic Acids Res. 32, 1792–1797.

Elgin, S.C.R., and Reuter, G. (2013). Position-effect variegation, heterochromatin formation, and gene silencing in Drosophila. Cold Spring Harb. Perspect. Biol. 5, a017780.

Enke, R.A., Dong, Z., and Bender, J. (2011). Small RNAs prevent transcription-coupled loss of histone H3 lysine 9 methylation in Arabidopsis thaliana. PLoS Genet. 7, 1–10.

Ghabrial, A., Ray, R.P., and Schupbach, T. (1998). okra and spindle-B encode components of the RAD52 DNA repair pathway and affect meiosis and patterning in Drosophila oogenesis. Genes Dev. 12, 27112723.

Gonzalez, I., Mateos-Langerak, J., Thomas, A., Cheutin, T., and Cavalli, G. (2014). Identification of regulators of the three-dimensional polycomb organization by a microscopy-based genome-wide RNAi screen. Mol. Cell 54, 485–499.

Gu, S.G., Pak, J., Guang, S., Maniar, J.M., Kennedy, S., and Fire, A. (2012). Amplification of siRNA in Caenorhabditis elegans generates a transgenerational sequence-targeted histone H3 lysine 9 methylation footprint. Nat Genet 44, 157–164.

Handu, M., Kaduskar, B., Ravindranathan, R., Soory, A., Giri, R., Elango, V.B., Gowda, H., and Ratnaparkhi, G.S. (2015). SUMO-Enriched Proteome for Drosophila Innate Immune Response. G3 (Bethesda). 5, 2137–2154.

Hari, K.L., Hari, K.L., Cook, K.R., Cook, K.R., Karpen, G.H., and Karpen, G.H. (2001). The Drosophila Su(var)2-10 locus regulates chromosome structure and function and encodes a member of the PIAS protein family. Genes Dev. 15, 1334–1348.

Holoch, D., and Moazed, D. (2015). RNA-mediated epigenetic regulation of gene expression. Nat Rev Genet 16, 71–84.

Ivanov, A. V., Peng, H., Yurchenko, V., Yap, K.L., Negorev, D.G., Schultz, D.C., Psulkowski, E., Fredericks, W.J., White, D.E., Maul, G.G., et al. (2007). PHD Domain-Mediated E3 Ligase Activity Directs Intramolecular Sumoylation of an Adjacent Bromodomain Required for Gene Silencing. Mol. Cell 28, 823–837.

Jacobs, S.A., Taverna, S.D., Zhang, Y., Briggs, S.D., Li, J., Eissenberg, J.C., Allis, Cd., and Khorasanizadeh, S. (2001). Specificity of the HP1 chromo domain for the methylated N-terminus of histone H3. EMBO J. 20, 5232–5241.

Jentsch, S., and Psakhye, I. (2013). Control of Nuclear Activities by Substrate-Selective and Protein-Group SUMOylation. Annu. Rev. Genet. 47, 167–186.

Johnson, E.S., and Gupta, A.A. (2001). An E3-like factor that promotes SUMO conjugation to the yeast septins. Cell 106, 735–744.

Kahyo, T., Nishida, T., and Yasuda, H. (2001). Involvement of PIAS1 in the sumoylation of tumor suppressor p53. Mol. Cell 8, 713–718.

Kaminker, J.S., Bergman, C.M., Kronmiller, B., Carlson, J., Svirskas, R., Patel, S., Frise, E., Wheeler, D.A., Lewis, S.E., Rubin, G.M., et al. (2002). The transposable elements of the Drosophila melanogaster euchromatin: a genomics perspective. Genome Biol. 3, RESEARCH0084.

Karolchik, D., Hinrichs, A.S., Furey, T.S., Roskin, K.M., Sugnet, C.W., Haussler, D., and Kent, W.J. (2004). The UCSC Table Browser data retrieval tool. Nucleic Acids Res. 32, D493–D496.

Kent, W.J., Zweig, A.S., Barber, G., Hinrichs, A.S., and Karolchik, D. (2010). BigWig and BigBed: enabling browsing of large distributed datasets. Bioinformatics 26, 2204–2207.

Klattenhoff, C., Bratu, D.P., McGinnis-Schultz, N., Koppetsch, B.S., Cook, H.A., and Theurkauf, W.E. (2007). Drosophila rasiRNA Pathway Mutations Disrupt Embryonic Axis Specification through Activation of an ATR/Chk2 DNA Damage Response. Dev. Cell 12, 45–55.

Koch, C.M., Honemann-Capito, M., Egger-Adam, D., and Wodarz, A. (2009). Windei, the Drosophila homolog of mAM/MCAF1, is an essential cofactor of the H3K9 methyl transferase dSETDB1/eggless in germ line development. PLoS Genet. 5, 1–15.

Kotaja, N., Karvonen, U., Janne, O.A., Jorma, J., and Ja, O.A. (2002). PIAS Proteins Modulate Transcription Factors by Functioning as SUMO-1 Ligases. Mol. Cell. Biol. 22, 5222–5234.

Krzywinski, M.I., Schein, J.E., Birol, I., Connors, J., Gascoyne, R., Horsman, D., Jones, S.J., and Marra, M.A. (2009). Circos: An information aesthetic for comparative genomics. Genome Res..

Kuramochi-Miyagawa, S., Watanabe, T., Gotoh, K., Totoki, Y., Toyoda, A., Ikawa, M., Asada, N., Kojima, K., Yamaguchi, Y., Ijiri, T.W., et al. (2008). DNA methylation of retrotransposon genes is regulated by Piwi family members MILI and MIWI2 in murine fetal testes. Genes Dev. 1, 908–917.

Lachner, M., O’Carroll, D., Rea, S., Mechtler, K., and Jenuwein, T. (2001). Methylation of histone H3 lysine 9 creates a binding site for HP1 proteins. Nature 410, 116–120.

Langmead, B., Trapnell, C., Pop, M., and Salzberg, S.L. (2009). Ultrafast and memory-efficient alignment of short DNA sequences to the human genome. Genome Biol. 10, R25.

LeThomas, A., Rogers, A.K., Webster, A., Marinov, G.K., Liao, S.E., Perkins, E.M., Hur, J.K., Aravin, A.A., and Toth, K.F. (2013). Piwi induces piRNA-guided transcriptional silencing and establishment of a repressive chromatin state. Genes Dev. 27, 390–399.

LeThomas, A., Marinov, G.K., and Aravin, A.A. (2014). A Transgenerational Process Defines piRNA Biogenesis in Drosophila virilis. Cell Rep. 8, 1–7.

Love, M.I., Huber, W., and Anders, S. (2014). Moderated estimation of fold change and dispersion for RNA-seq data with DESeq2. Genome Biol. 15, 550.

Maison, C., and Almouzni, G. (2004). HP1 and the dynamics of heterochromatin maintenance. Nat. Rev. Mol. Cell Biol. 5, 296–304.

Maison, C., Bailly, D., Roche, D., de Oca, R.M., Probst, A. V, Vassias, I., Dingli, F., Lombard, B., Loew, D., Quivy, J.-P., et al. (2011). SUMOylation promotes de novo targeting of HP1a to pericentric heterochromatin. Nat. Genet. 43, 220–227.

Maison, C., Bailly, D., Quivy, J.-P., and Almouzni, G. (2016). The methyltransferase Suv39h1 links the SUMO pathway to HP1a marking at pericentric heterochromatin. Nat. Commun. 7, 12224.

Manakov, S.A., Pezic, D., Marinov, G.K., Pastor, W.A., Sachidanandam, R., and Aravin, A.A. (2015). MIWI2 and MILI Have Differential Effects on piRNA Biogenesis and DNA Methylation. Cell Rep. 12, 1234–1243.

Meignin, C., and Davis, I. (2008). UAP56 RNA helicase is required for axis specification and cytoplasmic mRNA localization in Drosophila. Dev. Biol. 315, 89–98.

Mette, M.F., Aufsatz, W., van der Winden, J., Matzke, M.A., and Matzke, A.J. (2000). Transcriptional silencing and promoter methylation triggered by double-stranded RNA. EMBO J. 19, 5194–5201.

Mohr, S.E., and Boswell, R.E. (1999). Zimp encodes a homologue of mouse Miz1 and PIAS3 and is an essential gene in Drosophila melanogaster. Gene 229, 109–116.

Muerdter, F., Guzzardo, P.M., Gillis, J., Luo, Y., Yu, Y., Chen, C., Fekete, R., and Hannon, G.J. (2013). A genome-wide RNAi screen draws a genetic framework for transposon control and primary piRNA biogenesis in drosophila. Mol. Cell 50, 736–748.

Munarriz, E., Barcaroli, D., Stephanou, A., Townsend, P. a, Maisse, C., Terrinoni, A., Neale, M.H., Martin, S.J., Latchman, D.S., Knight, R. a, et al. (2004). PIAS-1 is a checkpoint regulator which affects exit from G1 and G2 by sumoylation of p73. Mol. Cell. Biol. 24, 10593–10610.

Nie, M., Xie, Y., Loo, J.A., and Courey, A.J. (2009). Genetic and proteomic evidence for roles of Drosophila SUMO in cell cycle control, Ras signaling, and early pattern formation. PLoS One 4.

Nishida, K.M., Iwasaki, Y.W., Murota, Y., Nagao, A., Mannen, T., Kato, Y., Siomi, H., and Siomi, M.C. (2015). Respective Functions of Two Distinct Siwi Complexes Assembled during PIWI-Interacting RNA Biogenesis in Bombyx Germ Cells. Cell Rep. 10, 193–203.

Ohtani, H., Iwasaki, Y.W., Shibuya, A., Siomi, H., and Siomi, M.C. (2013). DmGTSF1 is necessary for Piwi-piRISC-mediated transcriptional transposon silencing in the Drosophila ovary DmGTSF1 is necessary for Piwi-piRISC-mediated transcriptional transposon silencing in the Drosophila ovary. Genes Dev. 34283, 1656–1661.

Pezic, D., Manakov, S.A., Sachidanandam, R., Castro-diaz, N., Ecco, G., Coluccio, A., and Aravin, A.A. (2014). piRNA pathway targets active LINE1 elements to establish the repressive H3K9me3 mark in germ cells. Genes Dev. 28, 1410–1428.

Psakhye, I., and Jentsch, S. (2012). Protein group modification and synergy in the SUMO pathway as exemplified in DNA repair. Cell 151, 807–820.

Quinlan, A.R., and Hall, I.M. (2010). BEDTools: a flexible suite of utilities for comparing genomic features. Bioinformatics 26, 841–842.

Rahman, R., Chirn, G.-W., Kanodia, A., Sytnikova, Y.A., Brembs, B., Bergman, C.M., and Lau, N.C. (2015). Unique transposon landscapes are pervasive across Drosophila melanogaster genomes. Nucleic Acids Res. 43, 10655–10672.

Rangan, P., Malone, C.D., Navarro, C., Newbold, S.P., Hayes, P.S., Sachidanandam, R., Hannon, G.J., and Lehmann, R. (2011). piRNA production requires heterochromatin formation in drosophila. Curr. Biol. 21, 1373–1379.

Reindle, A., Belichenko, I., Bylebyl, G.R., Chen, X.L., Gandhi, N., and Johnson, E.S. (2006). Multiple domains in Siz SUMO ligases contribute to substrate selectivity. J. Cell Sci. 119, 4749–4757.

Reuter, G., and Wolff, I. (1981). Isolation of dominant suppressor mutations for position-effect variegation in Drosophila melanogaster. Mol. Gen. Genet. MGG 182, 516–519.

Rozhkov, N. V, Hammell, M., and Hannon, G.J. (2013). Multiple roles for Piwi in silencing Drosophila transposons. Genes Dev. 27, 400–412.

Ryu, T., Spatola, B., Delabaere, L., Bowlin, K., Hopp, H., Kunitake, R., Karpen, G.H., and Chiolo, I. (2015). Heterochromatic breaks move to the nuclear periphery to continue recombinational repair. Nat Cell Biol 17, 1401–1411.

Sachdev, S., Bruhn, L., Sieber, H., Pichler, A., Melchior, F., and Grosschedl, R. (2001). PIASy, a nuclear matrix-associated SUMO E3 ligase, represses LEF1 activity by sequestration into nuclear bodies. Genes Dev. 15, 3088–3103.

Schindelin, J., Arganda-Carreras, I., Frise, E., Kaynig, V., Longair, M., Pietzsch, T., Preibisch, S., Rueden, C., Saalfeld, S., Schmid, B., et al. (2012). Fiji: an open-source platform for biological-image analysis. Nat. Methods 9, 676–682.

Schmidt, D., and Muller, S. (2002). Members of the PIAS family act as SUMO ligases for c-Jun and p53 and repress p53 activity. Proc. Natl. Acad. Sci. 99, 2872–2877.

Seum, C., Reo, E., Peng, H., Rauscher, F.J., Spierer, P., and Bontron, S. (2007). Drosophila SETDB1 is required for chromosome 4 silencing. PLoS Genet. 3, 709–719.

Shin, Y., and Brangwynne, C.P. (2017). Liquid phase condensation in cell physiology and disease. Science 357, eaaf4382.

Shin, J.A., Eun, S.C., Hyun, S.K., Ho, J.C.Y., Watts, F.Z., Sang, D.P., and Jang, Y.K. (2005). SUMO modification is involved in the maintenance of heterochromatin stability in fission yeast. Mol. Cell 19, 817–828.

Sienski, G., Donertas, D., and Brennecke, J. (2012). Transcriptional silencing of transposons by Piwi and maelstrom and its impact on chromatin state and gene expression. Cell 151, 964–980.

Sienski, G., Batki, J., Senti, K.-A., Donertas, D., Tirian, L., Meixner, K., and Brennecke, J. (2015). Silencio/CG9754 connects the Piwi-piRNA complex to the cellular heterochromatin machinery. Genes Dev. 29, 1–14.

Stampfel, G., Kazmar, T., Frank, O., Wienerroither, S., Reiter, F., and Stark, A. (2015). Transcriptional regulators form diverse groups with context-dependent regulatory functions. Nature 528, 147–151.

Stielow, B., Sapetschnig, A., Kruger, I., Kunert, N., Brehm, A., Boutros, M., and Suske, G. (2008a). Identification of SUMO-Dependent Chromatin-Associated Transcriptional Repression Components by a Genome-wide RNAi Screen. Mol. Cell 29, 742–754.

Stielow, B., Sapetschnig, A., Wink, C., Kruger, I., and Suske, G. (2008b). SUMO-modified Sp3 represses transcription by provoking local heterochromatic gene silencing. EMBO Rep. 9, 899–906.

Takahashi, Y., Kahyo, T., Toh-E, A., Yasuda, H., and Kikuchi, Y. (2001). Yeast Ull1/Siz1 is a novel SUMO1/Smt3 ligase for septin components and functions as an adaptor between conjugating enzyme and substrates. J. Biol. Chem. 276, 48973–48977.

Thompson, P.J., Dulberg, V., Moon, K., and Foster, L.J. (2015). hnRNP K Coordinates Transcriptional Silencing by SETDB1 in Embryonic Stem Cells. 1–32.

Tzeng, T.-Y., Lee, C.-H., Chan, L.-W., and Shen, C.-K.J. (2007). Epigenetic regulation of the Drosophila chromosome 4 by the histone H3K9 methyltransferase dSETDB1. Proc. Natl. Acad. Sci. U. S. A. 104, 12691–12696.

Uchimura, Y., Ichimura, T., Uwada, J., Tachibana, T., Sugahara, S., Nakao, M., and Saitoh, H. (2006). Involvement of SUMO modification in MBD1-and MCAF1-mediated heterochromatin formation. J. Biol. Chem. 281, 23180–23190.

Verdel, A., Jia, S., Gerber, S., Sugiyama, T., Gygi, S., Grewal, S.I.S., and Moazed, D. (2004). RNAi-Mediated Targeting of Heterochromatin by the RITS Complex. Science (80-.). 303, 672 LP–676.

Volpe, T.A., Kidner, C., Hall, I.M., Teng, G., Grewal, S.I.S., and Martienssen, R.A. (2002). Regulation of Heterochromatic Silencing and Histone H3 Lysine-9 Methylation by RNAi. Science (80-.). 297, 1833 LP– 1837.

Waterhouse, R.M., Zdobnov, E.M., Tegenfeldt, F., Li, J., and Kriventseva, E. V. (2011). OrthoDB: the hierarchical catalog of eukaryotic orthologs in 2011. Nucleic Acids Res. 39, D283–D288.

Wolf, G., Greenberg, D., and Macfarlan, T.S. (2015). Spotting the enemy within: Targeted silencing of foreign DNA in mammalian genomes by the Kruppel-associated box zinc finger protein family. Mob. DNA 6, 17.

Yu, Y., Gu, J., Jin, Y., Luo, Y., Preall, J.B., Ma, J., Czech, B., and Hannon, G.J. (2015). Panoramix enforces piRNA-dependent cotranscriptional silencing. Science (80-.). 350, 339–342.

Zhao, Q., Xie, Y., Zheng, Y., Jiang, S., Liu, W., Mu, W., Liu, Z., Zhao, Y., Xue, Y., and Ren, J. (2014). GPS-SUMO: a tool for the prediction of sumoylation sites and SUMO-interaction motifs. Nucleic Acids Res. 42, W325–W330.

Zhu, J., Zhu, S., Guzzo, C.M., Ellis, N.A., Sung, K.S., Choi, C.Y., and Matunis, M.J. (2008). Small ubiquitin-related modifier (SUMO) binding determines substrate recognition and paralog-selective SUMO modification. J. Biol. Chem. 283, 29405–29415.

