## Supplemental_figures_and_table for "The SUMO ligase Su(var)2-10 links piRNA-guided target recognition to chromatin silencing"

### Suppl. Figure S1

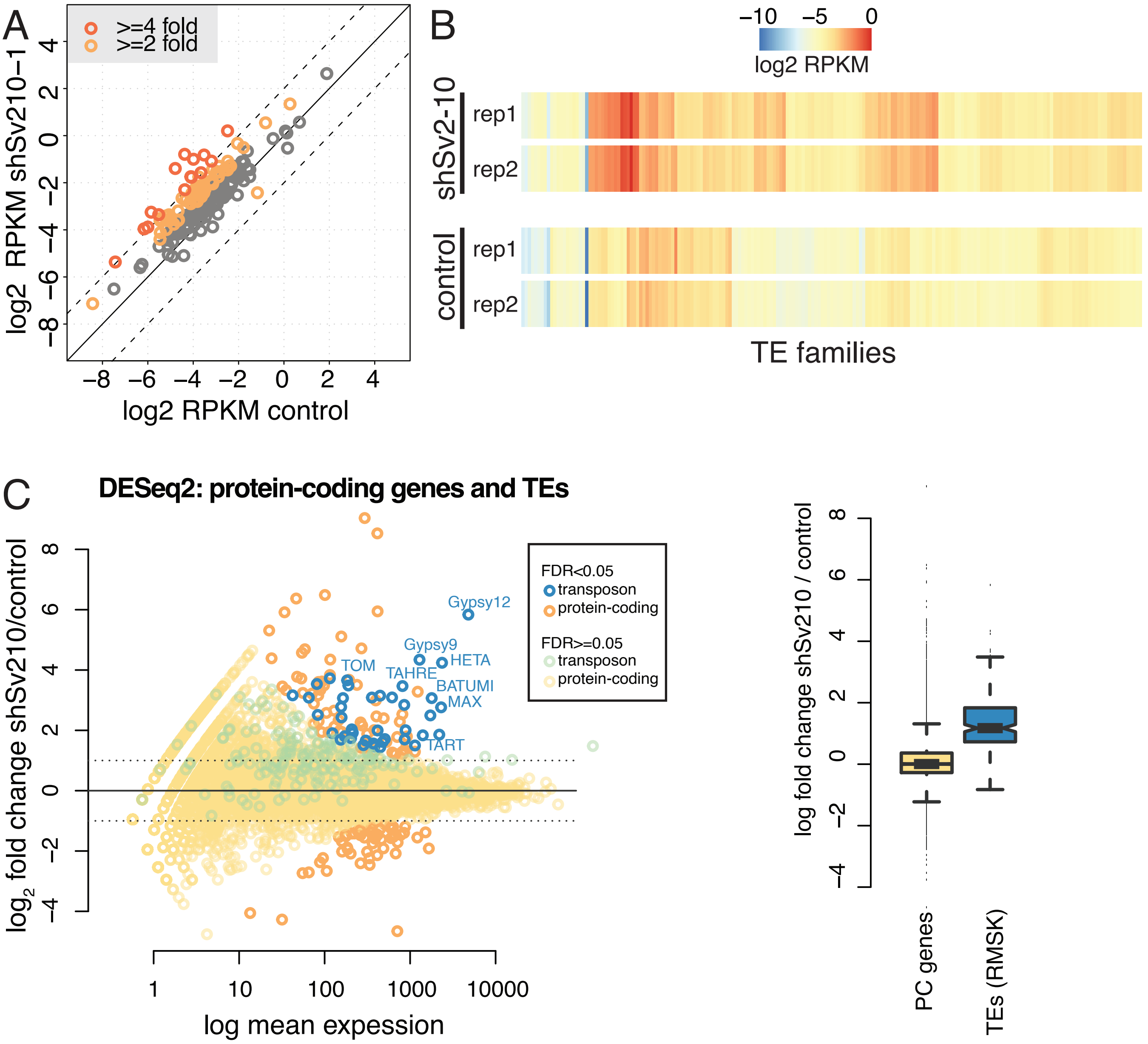

Suppl. Figure S2

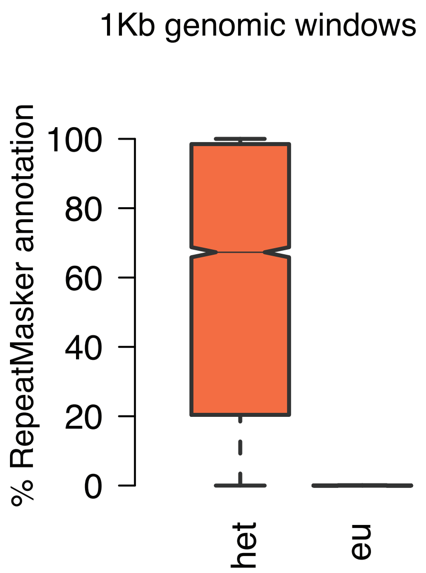

### Suppl. Figure S3

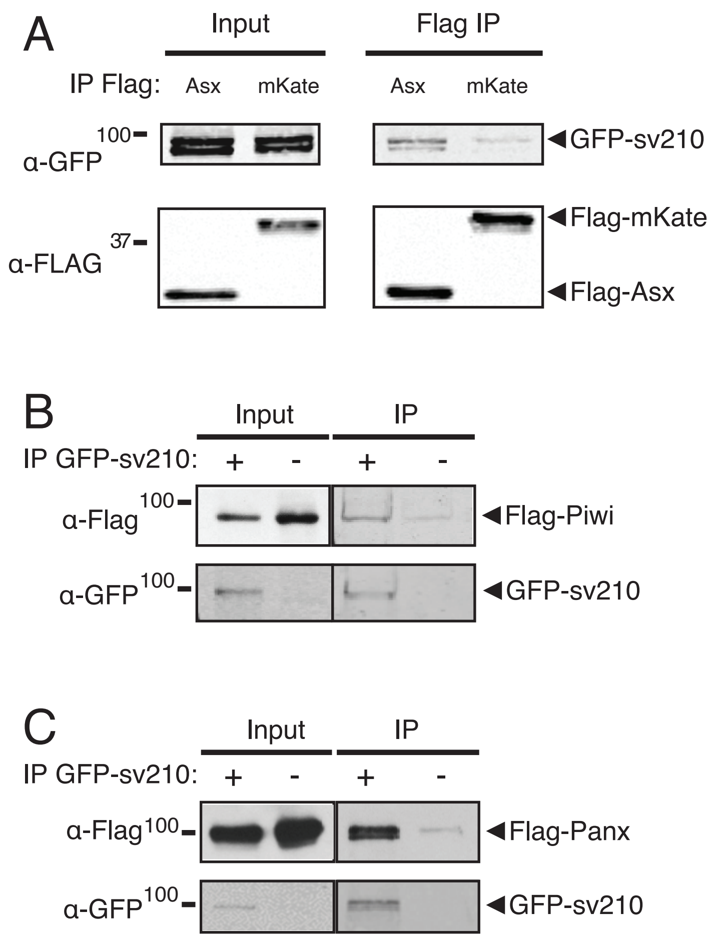

### Suppl. Figure S4

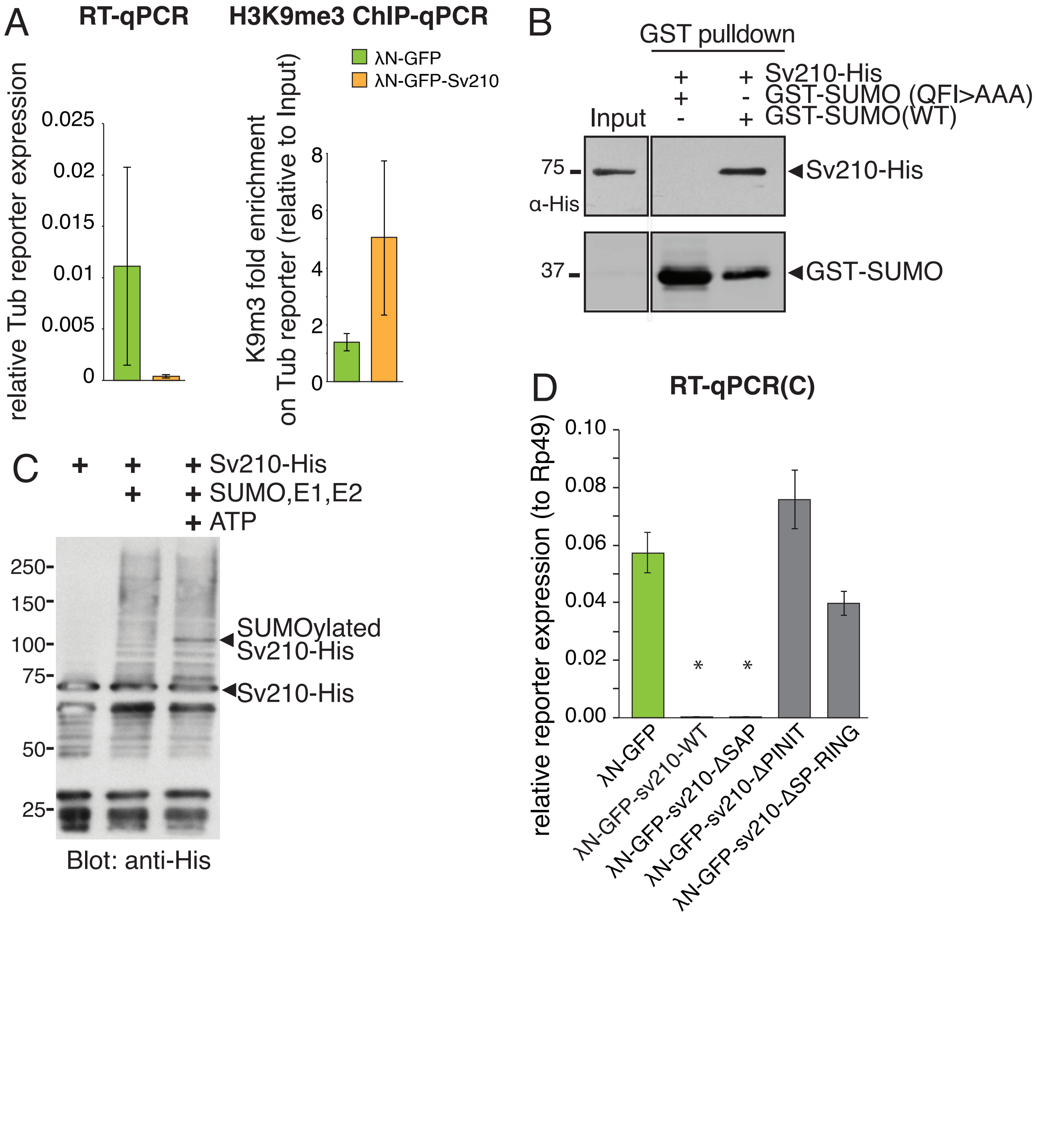

### Suppl. Figure S5

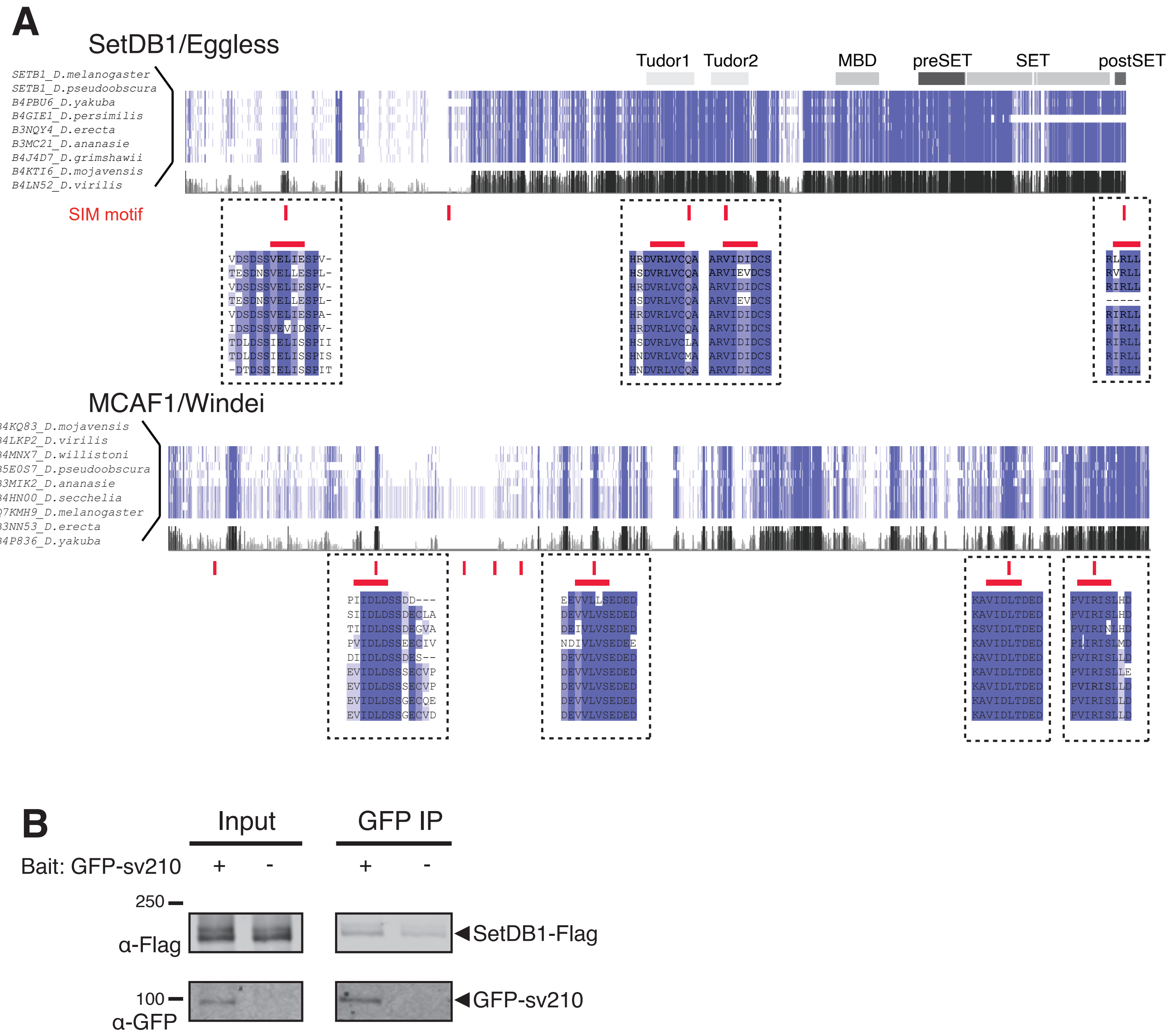

Supplementary Table S1. Primers list

| Name | Sequence |
|---|---|
| Su(var)2-10-F | CCAGCACAGGACGAACAGCCC |
| Su(var)2-10-R | CGTGGAACTGGCGACGGCTT |
| rp49-F | CCGCTTCAAGGGACAGTATCTG |
| rp49-R | ATCTCGCCGCAGTAAACGC |
| HetA-F | CGCGCGGAACCCATCTTCAGA |
| HetA-R | CGCCGCAGTCGTTTGGTGAGT |
| Blood-F | TGCCACAGTACCTGATTTCG |
| Blood-R | GATTCGCCTTTTACGTTTGC |
| ZAM-F | ACTTGACCTGGATACACTCACAAC |
| ZAM-R | GAGTATTACGGCGACTAGGGATAC |
| Ptip 3'UTR-F | CATGTGTGTTTCCGCCACAG |
| Ptip 3'UTR-R | TTCCCAGCTCGCGAAGAAAT |
| CG32138-F | CAGGATCTGCGCTACGACAT |
| CG32138-R | AATCGTCGGTCCAGCTCATC |
| sh-Asterix-F | CTAGCAGTCCAGTAGTTCGTGTTCATCAATAGTTATATTCAAGCA TATTGATGAACACGAACTACTGGGCG |
| sh-Asterix-R | AATTCGCCCAGTAGTTCGTGTTCATCAATATGCTTGAATATAACT ATTGATGAACACGAACTACTGGACTG |
| mKate2-F(A) | GTGACTGTGCGTTAGGTCCTG |
| mKate2-F(A) | TGAAGTGGTGGTTGTTCACGG |
| mKate2-F(B) | TCAGAGGGGTGAACTTCCCA |
| mKate2-R(B) | CTCCCAGCCGAGTGTTTTCT |
| mKate2-F(C) | GGCCGACAAAGAGACCTACG |
| mKate2-R(C) | CCAGTTTGCTAGGGAGGTCG |
| Tub-BoxB reporter-F | CTTCCTCCTCATCCACAGCG |
| Tub-BoxB reporter-R | ACTTGTGGCCGTTTACGTCG |
